# Bilallelic germline mutations in *MAD1L1* induce a novel syndrome of aneuploidy with high tumor susceptibility

**DOI:** 10.1101/2022.08.08.503198

**Authors:** Carolina Villarroya-Beltri, Ana Osorio, Raúl Torres-Ruiz, David Gómez-Sánchez, Marianna Trakala, Agustin Sánchez-Belmonte, Fátima Mercadillo, Borja Pitarch, Almudena Hernández-Núñez, Antonio Gómez-Caturla, Daniel Rueda, José Perea, Sandra Rodríguez-Perales, Marcos Malumbres, Miguel Urioste

**Author notes:** Corresponding authors (SRP), (MM), (MU).

## Abstract

Aneuploidy is a frequent feature of human tumors. Germline mutations leading to aneuploidy are very rare in humans, and their tumor-promoting properties are mostly unknown at the molecular level. We report here novel germline biallelic mutations in *MAD1L1*, the gene encoding the Spindle Assembly Checkpoint (SAC) protein MAD1, in a 36-year-old female with a dozen of neoplasias, including five malignant tumors. Functional studies in peripheral blood cells demonstrated lack of full-length protein and deficient SAC response, resulting in ∼30-40% of aneuploid cells as detected by cytogenetic and single-cell (sc) DNA analysis. scRNA-seq analysis of patient blood cells identified mitochondrial stress accompanied by systemic inflammation with enhanced interferon and NFkB signaling. The inference of chromosomal aberrations from scRNA-seq analysis detected inflammatory signals both in aneuploid and euploid cells, suggesting a non-cell autonomous response to aneuploidy. In addition to random aneuploidies, *MAD1L1* mutations resulted in specific clonal expansions of γδ T-cells with chromosome 18 gains and enhanced cytotoxic profile, as well as intermediate B-cells with chromosome 12 gains and transcriptomic signatures characteristic of chronic lymphocytic leukemia cells. These data point to *MAD1L1* mutations as the cause of a new aneuploidy syndrome with systemic inflammation and unprecedented tumor susceptibility.

## Introduction

Aneuploidy is a frequent finding in spontaneous abortions and a near-universal characteristic of tumor cells (*1, 2*). However, germline mutations leading to chromosomal instability are rare in humans and lead to clinical and genetically heterogeneous syndromes (*3*). The hallmark of these patients is the presence of mosaic aneuploidies, mainly trisomies and, more rarely, monosomies, and a variegated distribution of affected chromosomes in different cells and tissues throughout the body.

Human diseases characterized by chromosomal instability are typically associated to mutations in genes related to cell division. Mutations in centrosome regulators affect the majority of patients with primary microcephaly (*4*), and alteration of different cohesion subunits result in a variety of pathologies known as cohesinopathies (*5*). Aneuploidy, however, is only occasionally observed in these pathologies. On the other hand, mosaic variegated aneuploidy (MVA) typically correlates with mutations in components of the spindle assembly checkpoint (SAC), a multigene network that, together with other biological processes, controls the accurate chromosome segregation into daughter cells during cell division, and permits ordered cell cycle progression from metaphase to anaphase (*6, 7*). The SAC monitors the complete bipolar attachment of kinetochores on chromosomes to microtubules of the mitotic spindle, and delays anaphase entry in the presence of unattached kinetochores. This is achieved through the formation of the mitotic checkpoint complex (MCC), composed of BUBR1, MAD2, BUB3 and CDC20, and the subsequent inhibition of the Anaphase-promoting complex/Cyclosome (APC/C). SAC deficiency results in chromosome segregation in the presence of unattached chromosomes generating daughter cells with abnormal chromosome numbers. In patients, these alterations are associated with a heterogeneous and non-specific phenotype that typically involves pre- and postnatal growth retardation, eye and facial anomalies, variable developmental delay and intellectual disability (*8–11*).

Despite the widespread presence of aneuploidy in human tumors, to what extent chromosomal instability plays oncogenic or tumor suppressor roles is unclear (*1, 12*). Cancer predisposition is not a common feature in patients with microcephaly or cohesinopathies. MVA is associated with a higher increase in some childhood malignancies, such as rhabdomyosarcoma, Wilm’s tumor, and leukemia, although these neoplasias are only observed in a subset of MVA patients, and specific MVA-associated mutations have been suggested to impair tumor progression (*11, 13*). Common features of MVA patients are therefore intellectual disability and growth retardation, with or without microcephaly and tumors.

We describe here the first germline biallelic mutations in *MAD1L1* as a novel cause of aneuploidy in a patient with no intellectual disability and an unprecedented number of neoplasias. The *MAD1L1* product MAD1 (Mitotic arrest-deficient 1) is an essential component of the mitotic checkpoint originally identified as a protein required for mitotic arrest in the presence of mitotic poisons (*14, 15*). MAD1 recruits the MCC component MAD2 to unattached kinetochores thus promoting the formation of the MCC and APC/C inhibition (*16–18*). Lack of *Mad1l1* is lethal in the mouse during early development (*19*) and no germline pathogenic mutations in the *MAD1L1* gene have been described in humans. The *MAD1L1* patient display biallelic mutations that are present in her parents in a heterozygous manner. The patient developed 12 neoplasias including five malignant tumors before the age of 36. Cellular studies suggested defective SAC function associated to high levels of aneuploidy (30-40%) in peripheral blood cells. Single-cell transcriptomic studies allowed the first analysis of the molecular consequences of aneuploidy in a patient with germline chromosomal instability. These studies suggested the presence of a non-cell autonomous inflammatory response to aneuploidy, and identified small populations of cells with selected aneuploidies and specific functional or premalignant properties of physiological or clinical relevance.

## Results

### Identification of a patient with germline biallelic mutations in *MAD1L1*

The patient, a female, was born in 1986 after an uncomplicated at term pregnancy of a 26-year-old mother and a 30-year old father, both healthy and not consanguineous. Apgar was 9/10. Birthweight was 2,510 g (3rd<p<10th centile), with a length of 48 cm (10th<p<25th centile). Developmental delay and slightly psychomotor retardation were apparent during the first months of life (Table 1). Several café-au-lait spots were noted before the 6 months of life. At the age of 2 years, she had a stage III embryonal rhabdomyosarcoma of the left auditory canal that was treated with chemotherapy and radiotherapy. The impairment of her growth curve was interpreted as a consequence of the radiotherapy treatment and growth hormone therapy was indicated. In 2001, several bone masses suggesting enchondromatosis were seen in femur, humerus, and ulna. Also in 2001 a stage I-B clear cell cervical carcinoma with ectocervix and endocervix involvement was diagnosed, without evidences of HPV infection, and treated with hysterectomy, bilateral adnexectomy, brachytherapy and external radiotherapy. The patient had not been exposed to diethylstilbestrol (DES) in utero. In 2006. a pleomorphic adenoma of left parotid gland was surgically removed. One year later, she suffered a left mastoidectomy with left parotidectomy due to a low-grade fusiform cell sarcoma. During the period 2006-2010, several dysplastic nevi, a mammary lipoma and a pilomatrixoma were removed. In 2010, a hemitiroidectomy was performed by a multinodular goiter. In 2012, a polypectomy of one adenoma of the colon harboring an intramucosal adenocarcinoma, was carried out, and two years later, a pT3N0M0 rectal adenocarcinoma was resected. Another tubular adenoma was removed in the control colonoscopy in 2014.

**Table 1.**
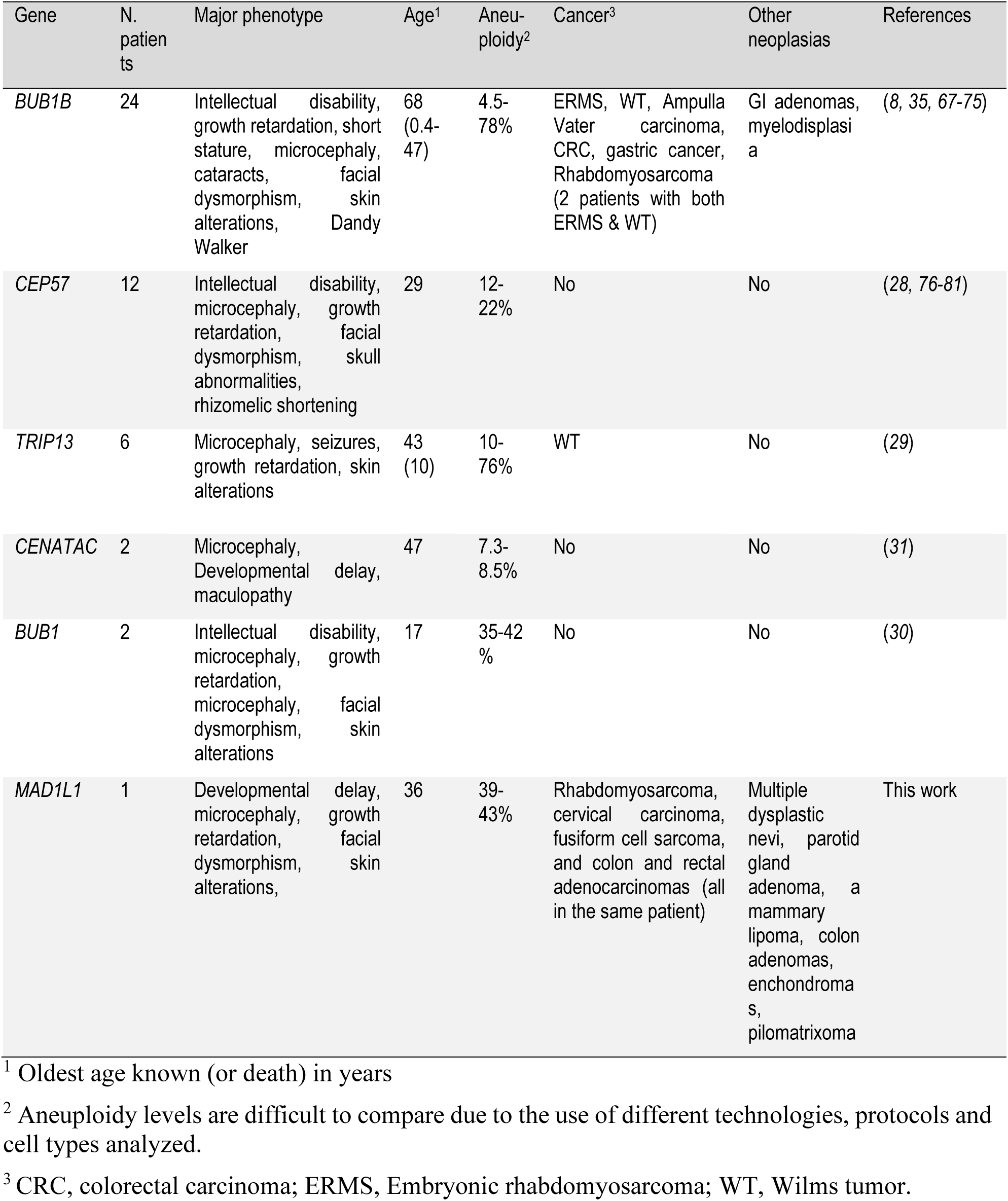
A comparison of phenotypes in patients with germline aneuploidy.

Physical examination in October 2017 revealed a weight of 44 kg (3rd centile), height 146 cm (p<3) and occipitofrontal circumference 50 (p<3). She had bitemporal narrowing, deeply set eyes, a midface hypoplasia, micrognathia, low-set and posteriorly rotated ears, nystagmus, bilateral pes cavus and bilateral hammer toes. She showed very striking cutaneous features with multiple widespread hyperpigmented lesions of variable tone. Hyperpigmented lesions in the anterior part of the trunk had a quadrant/blaschkoid wide band distribution affecting both sides of the body, and millimeter clear pigmented macular lesions on the back of the hands and feet, right axillae and right iliac fossa. Alternating with pigmented areas, she had patchy areas of hypopigmented and healthy skin. The hypopigmentation was blaschkoid in narrow band in both upper extremities and the left pretibial region, and with more erratic/phylloides distribution in the rest of the body. She showed hyperkeratotic plaques with furfuraceous desquamation on her knees and compact hyperkeratosis in the soles of the feet in support areas (Fig. S1A-B).

We performed biopsies of the hyper, hypopigmented and healthy skin and the hyperkeratotic areas of the knee. Optical miscroscopy with hematoxilin-eosin staining, showed basal pigmentation in the hyperpigmented and healthy skin, more striking in the hyperpigmented skin. In the samples of healthy and hypopigmented skin, scattered melanocytes were observed. Immunohistochemistry with Melan-A technique evidenced an increase in basal melanocytes in hyperpigmented and normal skin, with virtually no pigment in the basal hypopigmented skin. The biopsy of the hyperkeratotic zone showed a subacute spongiosis dermatitis, with irregular descending hyperplasia of the epidermis with foci of spongiosis with vesiculation and slight lymphocyte exocytosis, and superficial perivascular lymphocytic infiltrate in the dermis. The radiographic study of the upper and lower extremities revealed the existence of lesions suggestive of enchondromas in the left femur and in the right arm (Fig. S1C-D). There were not evidences of mental retardation, hypotonia, seizures, progeroid traits or immunodeficiency.

We initially ruled out mutations in *MLH1, MSH2, MSH6, PMS2, NF1* and *TP53* by focused DNA sequencing to discard alterations in genes related to Constitutional Mismatch Repair Deficiency, Neurofibromatosis and Li-Fraumeni syndromes. Furthermore, the colorectal and endometrial tumors of the patient were microsatellite stable, and the tumor immunohistochemistry analysis results in conserved expression of the four mismatch repair proteins. Whole Exome Sequencing (WES) of peripheral blood samples from the proband identified two different stop-gained mutations in *MAD1L1*: c.196C>T; p.Q66* in exon 4, and c.1882G>T; p.E628* in exon 18 (Fig. 1A). The first one, affecting most of the protein, has been described in a European individual (ExAC; MAF of 8.304e-06). The second one is not present in ExAC, Gnomad, Cosmic or The Cancer Genome Database. This mutation prevents the expression of the globular head of the C-terminal domain (CTD; Fig. 1B,C), a region critical for binding to MAD2 or CDC20, and for SAC activity (*20, 21*). By Sanger analysis, we determined that the proband’s mother and other maternal relatives carry the first mutation, while the second one had a paternal origin (Fig. 1D and Fig. S1E).

**Fig. 1.**
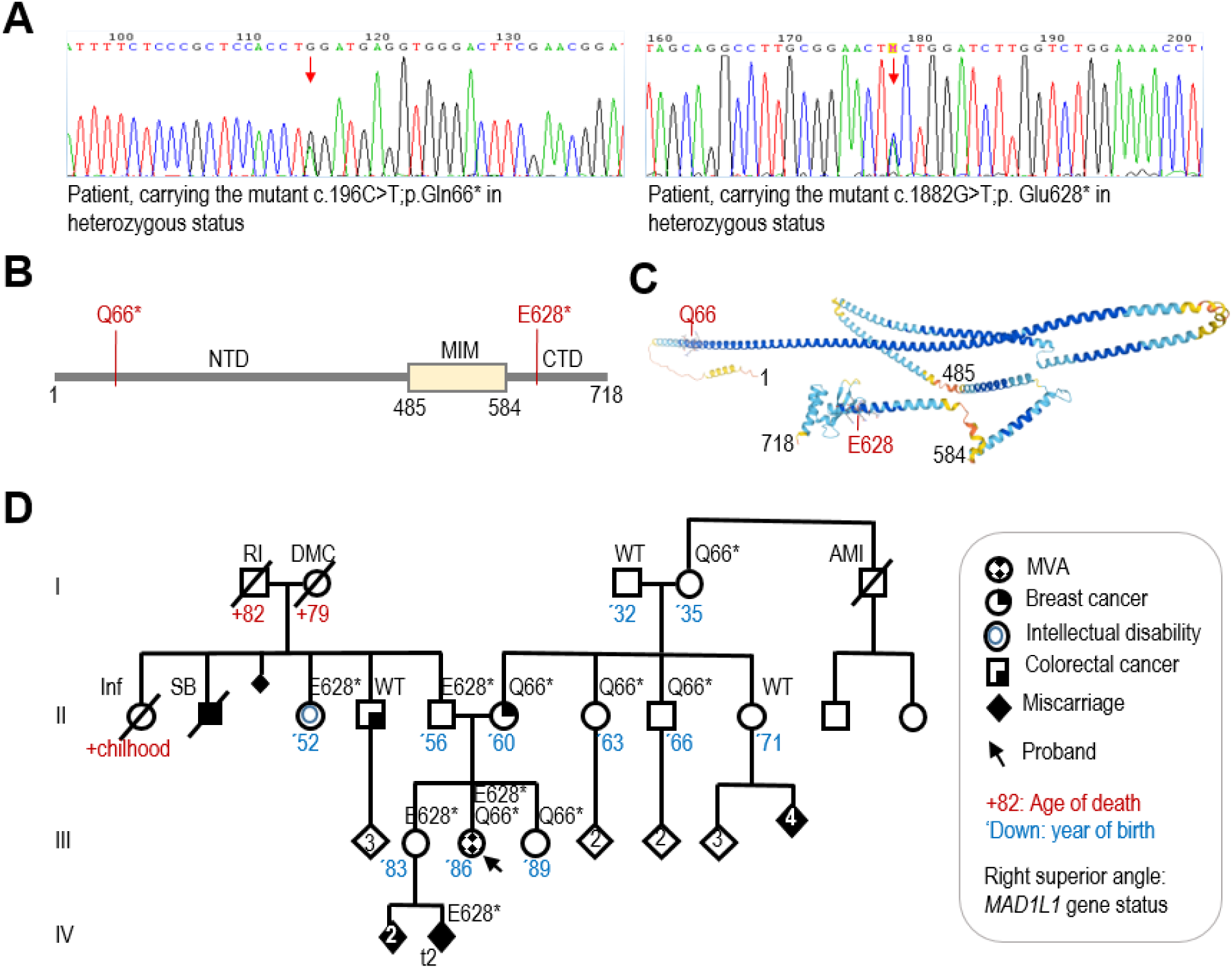
Bilallelic loss-of-function mutations in *MAD1L1*. **A**, Chromograms showing the two heterozygous mutations in the MAD1L1 gene in the proband. **B**, Schematic representation of the MAD1 protein and the mutations found in the proband. NTD, N-terminal domain; MIM, MAD2-interaction motif; CTD, C-terminal domain. **C**, Predicted structure of MAD1 (AlphaFold) showing critical residues limiting the different domains and the position of the Q66 and E628 residues mutated. **D**, Pedigree of the family with the status of *MAD1L1* gene shown on the right superior angle of each individual. AMI, Acute myocardial infarctation; DMC, Diabetes Mellitus complications; Inf, Infection; RI, renal insufficiency; SB, stillbirth; t2, trisomy chromosome 2.

### *MAD1L1* mutations induce germline and tumor aneuploidy

Metaphase spreads in peripheral blood mononuclear cells (PBMCs) identified 39% of aneuploid metaphases in the proband (Fig. 2A). The number of chromosomal gains found in the parents was low (0.7% in the father and 1.5% in the mother) and these alterations were not found in the sisters or healthy controls. Analysis of specific aberrations in chromosomes 7, 8, 20 and X using fluorescent in situ hybridization (FISH) confirmed that copy number gains in these chromosomes were more frequent than losses (Fig. 2B and Fig. S2A). Further studies using shallow whole genome sequencing in single cells identified whole-chromosome aneuploidies in 10 out of 23 (43.5%) proband cells but not in controls (Fig. 2C and Fig. S2B). Trisomy of chromosome 21 was the most frequent alteration in addition to recurrent gains of chromosomes 12 and 18. Monosomies were not observed with this technique and del(20p) was observed in one of the cells (Fig. 2C). CGH analysis of normal and pathological tissues identified chromosomal gains in most pathological conditions whereas losses were only observed in colorectal carcinoma (CRC). The progressive acquisition of CNV is evident from tubular adenoma, with gain of chromosome 7 as unique alteration, to CRC with many additional alterations (Fig. 2D).

**Fig. 2.**
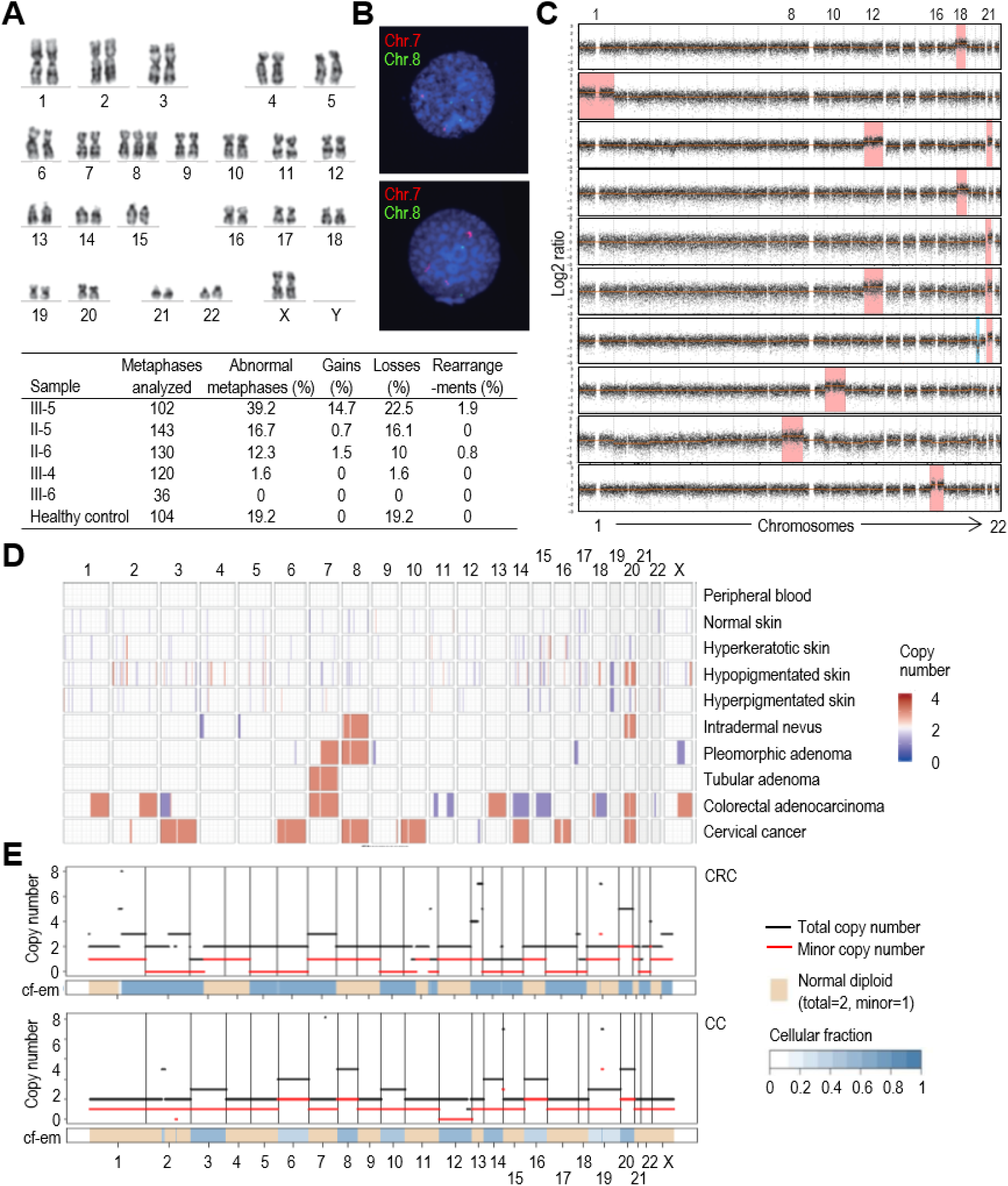
Analysis of aneuploidy. **A**, Representative example of the analysis of metaphase spreads from peripheral blood cells. The image shows an example of trisomy of chromosome 8 in the proband. The table in the bottom of the panel shows the summary of cytogenetic analysis of G-banding karyotype of family members proband (III-5), father (II-5), mother (II-6), older sister (III-4), younger sister (III-6) and a 46,XX healthy control. **B**, Examples of two-color FISH with centromeric chromosomes 7 (red) and 8 (green) DNA probes. Representative chromosome 8 trisomic nucleus with two signals for chromosome 7 and three signals for chromosome 8 (upper nucleus); and representative chromosome 7 trisomic nucleus with three signals for chromosome 7 and two signals for chromosome 8 (lower nucleus). See Fig. S2A for quantitative data. **C**. Detection of aneuploidies using single-cell DNA sequencing. Nine aneuploid cells from the proband are shown. Red and green indicates significant increase or decrease in copy number, respectively. **D**, CNV plot calculated from aCGH data of ten III-5 samples including peripheral blood, skin, and different types of cancer. (blue=copy number loss; red=copy number gain). **e**, FACET view representing copy number inferred from WES data in different neoplasias of the proband. CRC, rectal adenocarcinoma; CC, cervical carcinoma.

We analyzed genomic changes in neoplasias developed by the proband by WES. FACETS (Fraction and Allele-Specific Copy Number Estimates from Tumor Sequencing, (*22*)) analysis suggested that, in agreement with the previous observations, gains of one or several complete chromosomes were relatively frequent, while losses of complete chromosomes were observed only in CRC (Fig. 2E and Fig. S3A). Total copy number changes were higher in more severe neoplasias (CRC and CC), being remarkable the high proportion of uniparental alterations (partial or complete disomies, trisomies or tetrasomies) in the CRC. However, these uniparental alterations were infrequent in CC, only involving chromosome 12. The somatic variants found in the proband’s neoplastic tissues corresponded to canonical tumorigenesis pathways in these pathologies, including mutations in *APC* (p.R1255X), *TP53* (p.V25F), *RET* (p.S922Y), *KRAS* (p.G12A) in ICRC and CRC (Fig. S3B). None of these mutations was identified in TA, neither in other tissues sequenced. *CTNNB1* mutations and a frameshift mutation in *POLE* were identified in PA. Missense mutations in *RECQL4* and *FAT4* were the most common alterations in CC (Fig. S3B). CGH analysis of the third abortion in the older sister (III-4) identified a trisomy in chromosome 2 (Fig. S3C). CGH analysis of the ductal breast cancer of the proband’s mother showed mainly partial chromosome gains and losses, with reduced whole chromosome copy number alterations, as usually observed in sporadic breast cancer (Fig. S3D,E). WES analysis of this tumor identified the heterozygous *MAD1L1* p.(Q66*) mutation in addition to new mutations in *TP53* and *PIK3CA* (Fig. S3B).

### Mutations in *MAD1L1* result in defective SAC function

*MAD1L1* mutations resulted in lack of full-length MAD1 protein in peripheral lymphocytes from the proband, and reduced levels in lymphocytes from the heterozygous parents (Fig. 3A). Unexpectedly, mutant lymphocytes proliferated normally after mitogenic stimuli (Fig. 3B) and no obvious defects were observed in mitotic figures from early-passage cultures. However, proband and to a lesser extent mother cells displayed reduced load of BUBR1, the protein encoded by the *BUB1B* gene mutated in MVA1, in kinetochores lacking attached microtubules in the presence of taxol (Fig. 3C). In addition, the taxol-induced mitotic arrest was deficient in lymphocytes from the proband (Fig. 3D) and, accordingly, a larger amount of aberrant nuclei with abnormal shape and lobuli, as well as interphasic cells with micronuclei, and increased nuclear volume were observed, suggesting premature exit and/or abnormal recovery from taxol in the presence of a missfunctional SAC (Fig. 3E).

**Fig. 3.**
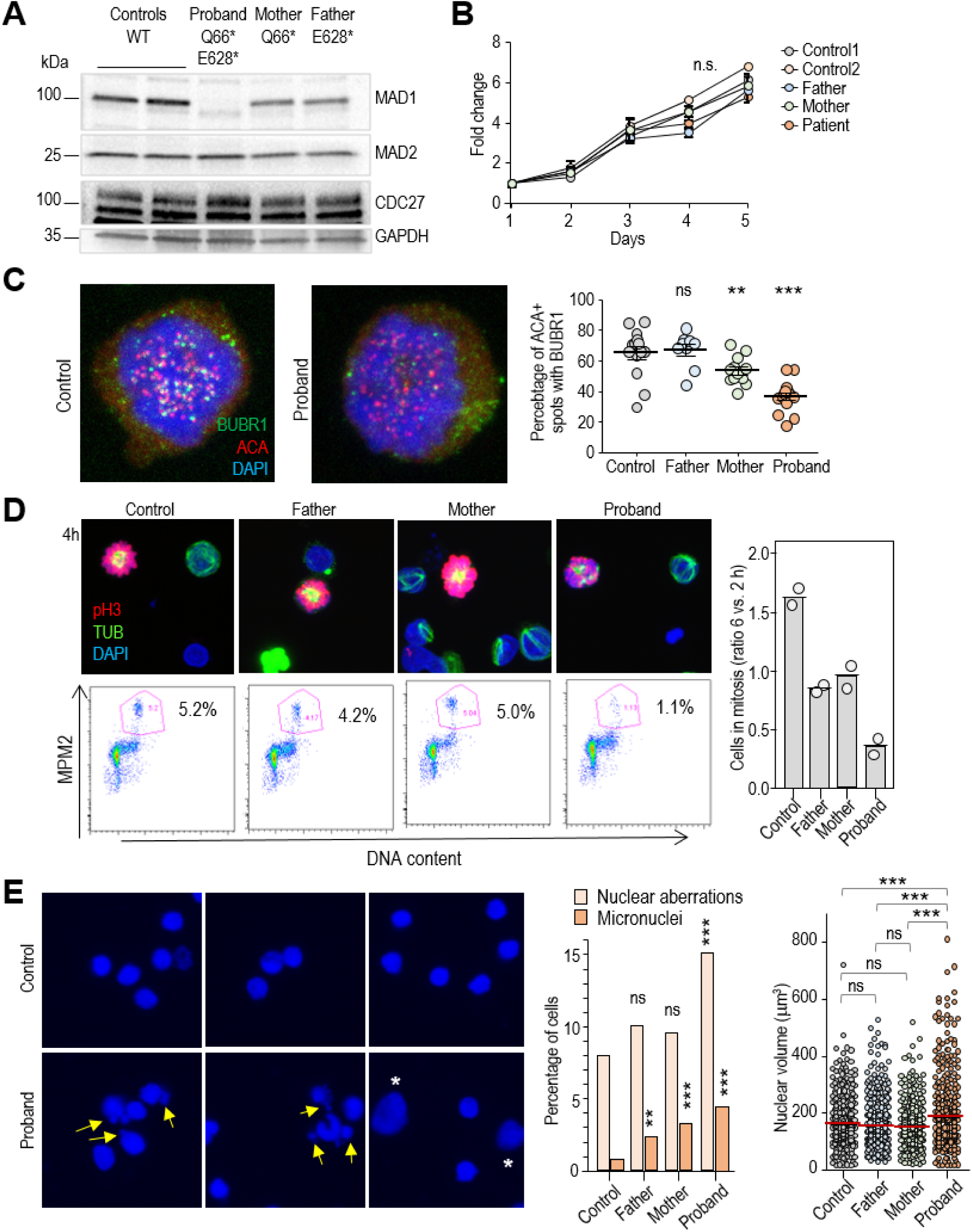
Defective chromosome segregation in biallelic *MAD1L1* mutant lymphocytes. **A**, Immunoblot showing lack of normal MAD1 proteins in the proband. The status of the *MAD1L1* gene is indicated for each sample. **B**, Proliferation of lymphocytes isolated from the peripheral blood of the indicated individuals. Data are mean ± S.D. (n=3 replicates). **C**, Immunofluorescence of BUBR1 and anti-centromeric antigens (ACA) in control and proband lymphocytes treated with taxol. The percentage of ACA+ centromeres with nearby BUBR1 signals is show to the right. **D**, Representative micrographs (top) and FACS analysis (bottom) of lymphocytes from the indicated individuals after 4 h in the presence of taxol. Top panels show immunofluorescence for phosphorylated histone H3 (pH3, red), α-tubulin (green) and DAPI staining for DNA (blue). Bottom panels show the percentage of cells positive for the mitotic antigens MPM2. The percentage of cells in mitosis after 6 h in taxol normalized versus the number of cells that entered mitosis at 2 h is show in the bar plot to the right (n=2 independent experiments). **E**, Representative example of interphase cells after treatment with taxol showing aberrant cells (abnormal shape and fragmented lobuli) and micronuclei (arrows), as well as cells large nuclei (asterisks) in mutant lymphocytes. The percentage of nuclear aberrations and cells with micronuclei, as well as quatification of nuclear volume is shown in the plots to the right. In C,E, ns, not significant; **, p<0.01; ***, p<0.001, one-way ANOVA with Bonferrini correction (C and E, nuclear volume) or ChiSquare, two-coiled Fisher exact test (E, nuclear aberrations and micronuclei).

### A systemic inflammation response to aneuploidy

To understand the consequences of aneuploidy in vivo, we sequenced a total of 33,604 individual PBMCs from the patient, both heterozygous parent and two healthy donors (Fig. 4A and Fig. S4A) and we assigned each cell, using available algorithms for PBMC classification (*23*), to 8 (L1), 28 (L2) or 48 (L3) different cell types (Fig. 4B and Fig. S4B,C). The relative abundance of the different cell types in all samples suggested the expansion of T-cells expressing the delta and gamma chains of the T-cell receptor (*TRD* and *TRG* genes; γδT-cells) and, to a lesser extent, B-cells expressing the kappa light chain (*IGKC*) in the proband (Fig. 4C-E and Fig. S4D).

**Fig. 4.**
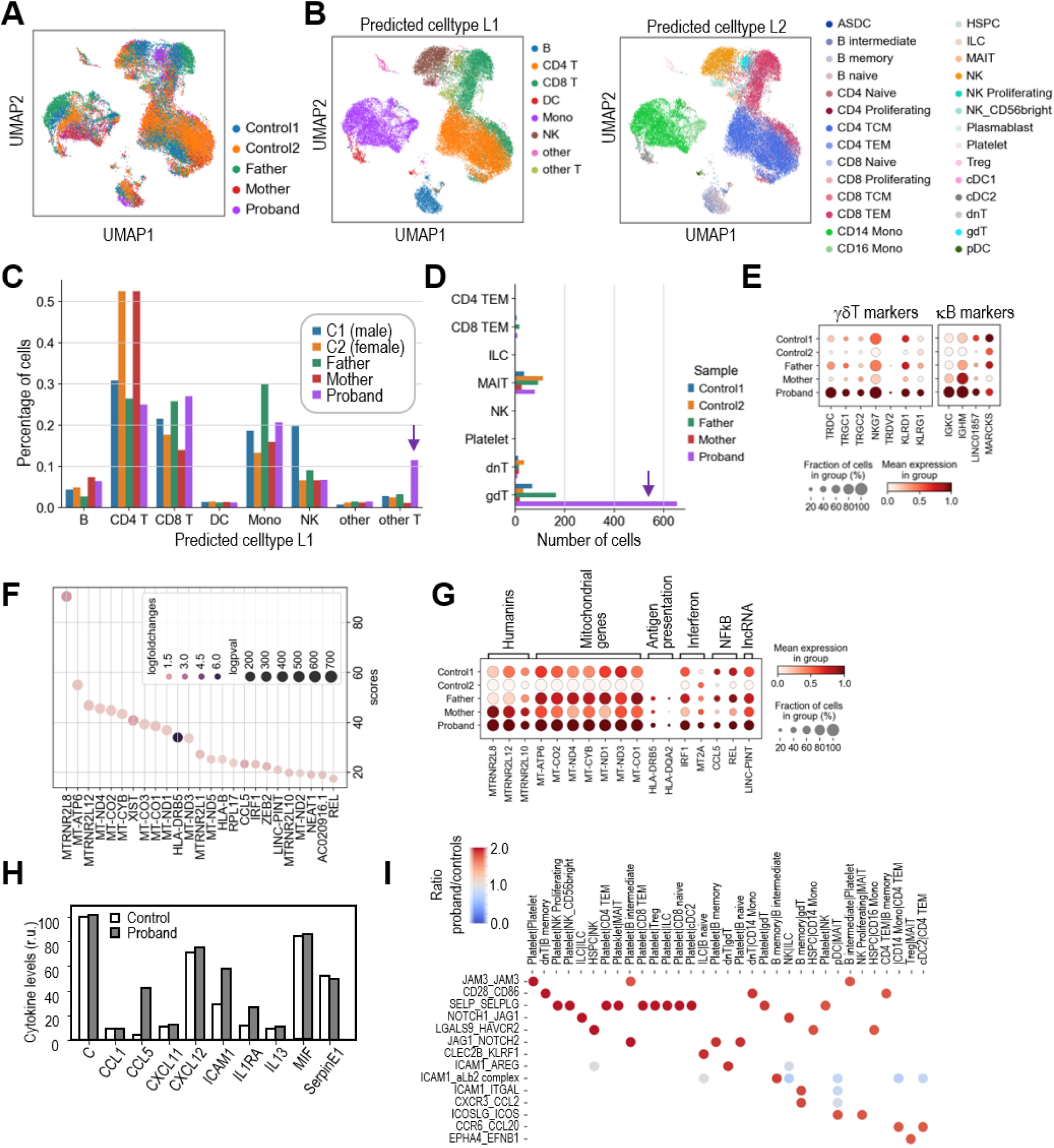
Transcriptomic analysis of peripheral blood cells with biallelic *MAD1L1* mutations. **A**, Uniform Manifold Approximation and Projection (UMAP) of cells from the indicated samples. **B**, UMAP distribution of cell types (L1 and L2 classifications following the Azimuth algorithms). **C**, Percentage of L1 cell types in the indicated samples. **D**, Number of cells in the specific cell types included in the L1 “other T” classification showing an overrepresentation of γδT-cells in the proband. **E**, Expression of specific markers of γδT-cells and B-cells expressing the kappa chain. **F**, List and scores (Wilcoxon rank sum test) of the top upregulated genes in proband versus control peripheral blood mononuclear cells (PBMCs). **G**, Aggregated expression of the set of top genes upregulated in proband cells in the different individuals analyzed. **H**, Relative levels of a panel of cytokines in the plasma of controls and the proband. **I**, Top ligand-receptor interaction pairs over-represented in the proband compared to the controls. The scale represents the ratio between the significant means for each ligand-receptor interaction pair (y axis) and pair of interacting cell types (x axis) in the proband vs. controls.

Analysis of the 25-top genes upregulated in proband PBMCs versus control cells identified multiple mitochondrial genes (*MT-ATP6, MT-ND4, MT-CO2, MT-CYB,* etc.), as well as several members of the humanin family, being the humanin-like MTRNR2L8 the top-scoring upregulated gene in proband cells (Fig. 4F,G). Interestingly, humanins have been shown to be released to the extracellular milieu upon mitochondrial stress and to act as mitokynes (mitochondrial cytokines) with cytoprotective effects (*24*). Among the most upregulated genes we could also detect the chemokine *CCL5* and members of the interferon (*IRF1, MT2A*) and NFkB (*REL*) signaling pathways. In accordance, CCL5 upregulation was also detected in the plasma of the proband together with ICAM1(Fig. 4H). A specific analysis of ligand-receptor interactions (*25*) suggested increased signals involved in platelet activation as well as molecules involved in antigen presentation and interferon signaling, NK and γδ T-cell activation, recruitment of leukocytes to circulation sites of inflammation and other general inflammatory signals (Fig. 4I). The strongest differences between proband and control ligand-receptor interactions were found in molecules expressed in platelets, such as the Junctional Adhesion Molecule 3 (JAM3) or the Selectin P (SELP). Other ligand-receptor interaction pairs overrepresented in proband cells included molecules involved in antigen presentation and interferon signaling, NK and γδ T-cell activation, recruitment of leukocytes to circulation sites of inflammation and other general inflammatory signals (Fig. 4I).

Pathway enrichment analysis suggested significant downregulation of pathways related to protein synthesis (ribosome biogenesis, translation and RNA processing), along downregulation of oxidative phosphorylation (OXPHOS) and MYC targets (Fig. 5A and Fig. S5A,B). On the other hand, proband cells displayed enhanced expression of genes related to inflammatory response, including interferon and NFκB signaling pathways. Whereas downregulation of ribosome and OXPHOS was shared by most cell types, inflammatory pathways were most evident in T-cells (Fig. 5A and Fig. S5A-C). Pathways related with antigen presentation were also significantly upregulated in proband’s cells, with class-II MHC molecules between the most upregulated genes in the proband, especially in professional antigen presenting cells (DCs, monocytes and B cells) and in the expanded population of γδ T cells (Fig. 5A and Fig. S5C,D).

**Fig. 5.**
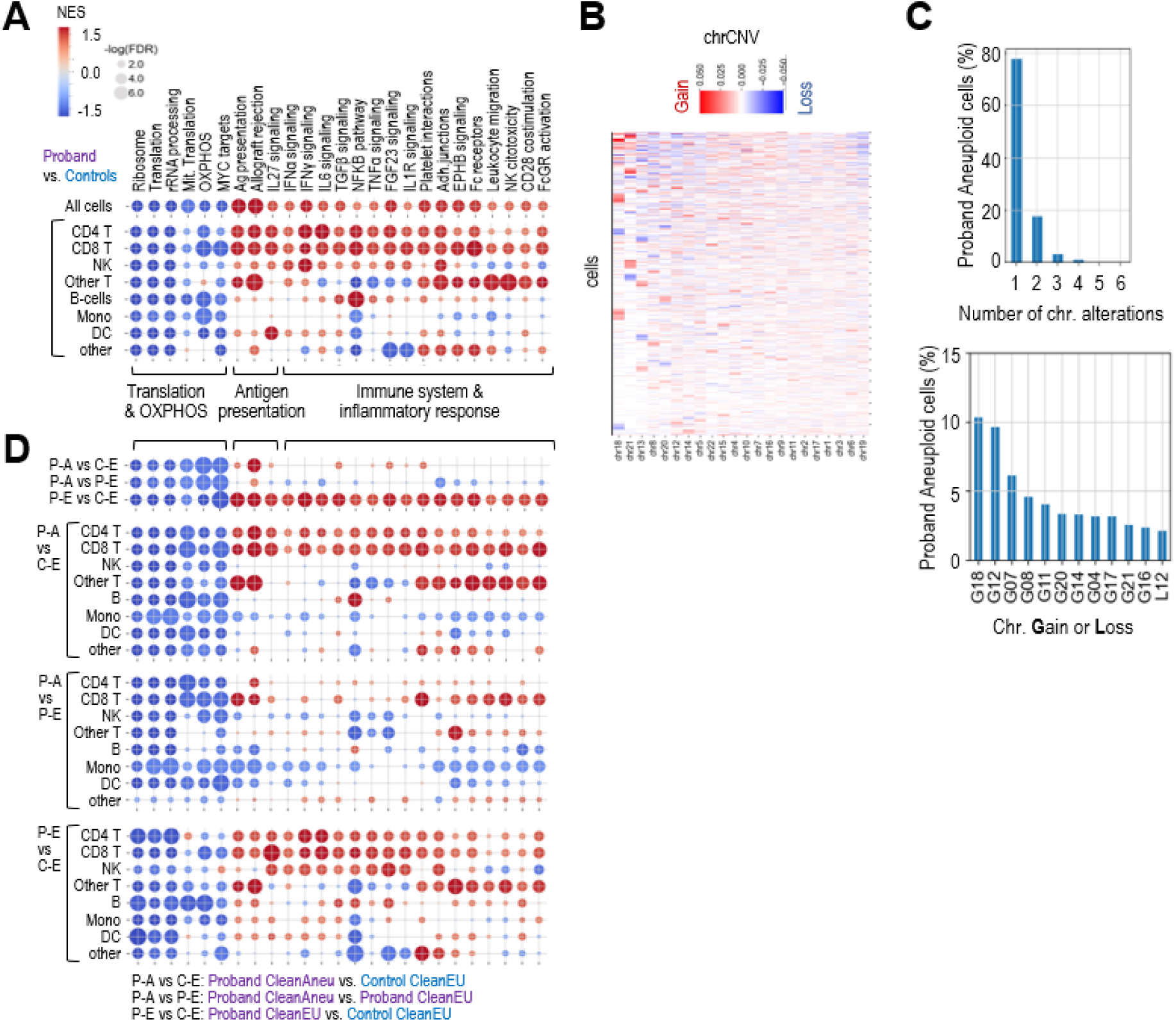
Pathway analysis of aneuploid and euploid *MAD1L1* mutant cells. **A**, Gene Set Enrichment Analysis (GSEA) used pre-ranked differential expression of genes in the indicated comparisons. The list of pathways represents a selection of differentially regulated pathways in the different conditions. **B**, Plot showing the average CNV score per chromosome (chrCNV score). Chromosomes are ordered by the number of extreme values in the chrCNV score; i.e. from chromosomes with higher monosomy or trisomy scores (left) to chromosomes with higher representation or disomies (right). **C**, Distribution of cells with numerical alterations in one or more chromosomes in the proband (top), and percentage of cells with the indicated aneuploidies in the proband (bottom). G, gain; L, loss of the indicated chromosomes. **D**, Deregulation of the selected pathways in proband aneuploid cells (P-A) vs. control euploid (C-E) cells, proband aneuploid cells vs. proband euploid cells (P-A vs. P-E), and proband euploid cells versus control euploid cells (P-E vs. C-E). Scale bar as in panel A.

To estimate chromosomal gains and losses, we made use of the inferCNV algorithm (*26*), which infers DNA copy number changes from the gene expression data, using matched control cell types for normalization (see Materials and Methods for details). The estimation of aneuploidies from scRNA-seq data using suggested a wide spectrum of chromosomal changes in different cell types (Fig. 5B and Fig. S6A,B). The lowest ratios of aneuploidy were found in progenitor cells, whereas 67.9% of intermediate (or transitional) B-cells were aneuploid in the proband (Fig. S6C). Most aneuploid cells contained a single chromosomal gain or loss (77.8%), and gains of chromosomes 18 and 12 were the most abundant alterations (10.3% and 9.7% of all aberrations; Fig. 5C).

We next compared the transcriptomic profile of aneuploid cells by using a more stringent threshold to define “clean euploid” (CleanEU) and “clean aneuploid” (CleanAneu) cells (Fig. S6B,C), thus avoiding the high noise in the InferCNV approach. Only cells with at least one mono- or trisomy with the stringent thresholds were considered as CleanAneu (13.6% in the proband vs. 4.0% in controls), and only cells with all disomic chromosomes using these stringent conditions were classified as CleanEU. Interestingly, analysis of the 25-gene signature overexpressed in the proband suggested that these genes were significantly overexpressed in aneuploid cells, but also to a lesser extent in CleanEU cells from the proband (Fig. S6D). Pathway analysis detected the downregulation of translation and OXPHOS, and upregulation of antigen presentation-related pathways in CleanAneu cells from the proband when compared to CleanEU cells from either the proband or controls. However, aneuploid and euploid cells from the proband did not show differences in immune pathways, whereas this inflammatory response was evident when comparing CleanEU cells from the proband with CleanEU cells from the controls (Fig. 5D). Together, all these data suggest that aneuploidy induces a cell-autonomous response characterized by increased antigen presentation and downregulation of translation and OXPHOS, and a systemic non-cell-autonomous inflammatory response.

### Clonal expansion of specific B- and T-cell subsets

Whereas random aneuploidies affected most cell types, we observed a preference for chromosome 18 gain in γδ T cells, and chromosome 12 gain in intermediate B-cells expressing the κ light chain subunit, and to a lesser extent the λ subunit, frequently accompanied of concomitant gain of chromosome 21 (Fig. 6A,B and Fig. S7A,B). Both intermediate B cells and γδ T cells were expanded in the proband (Fig, 6B and Fig. S4D), suggesting clonal or subclonal expansion of aneuploid cells.

**Fig. 6.**
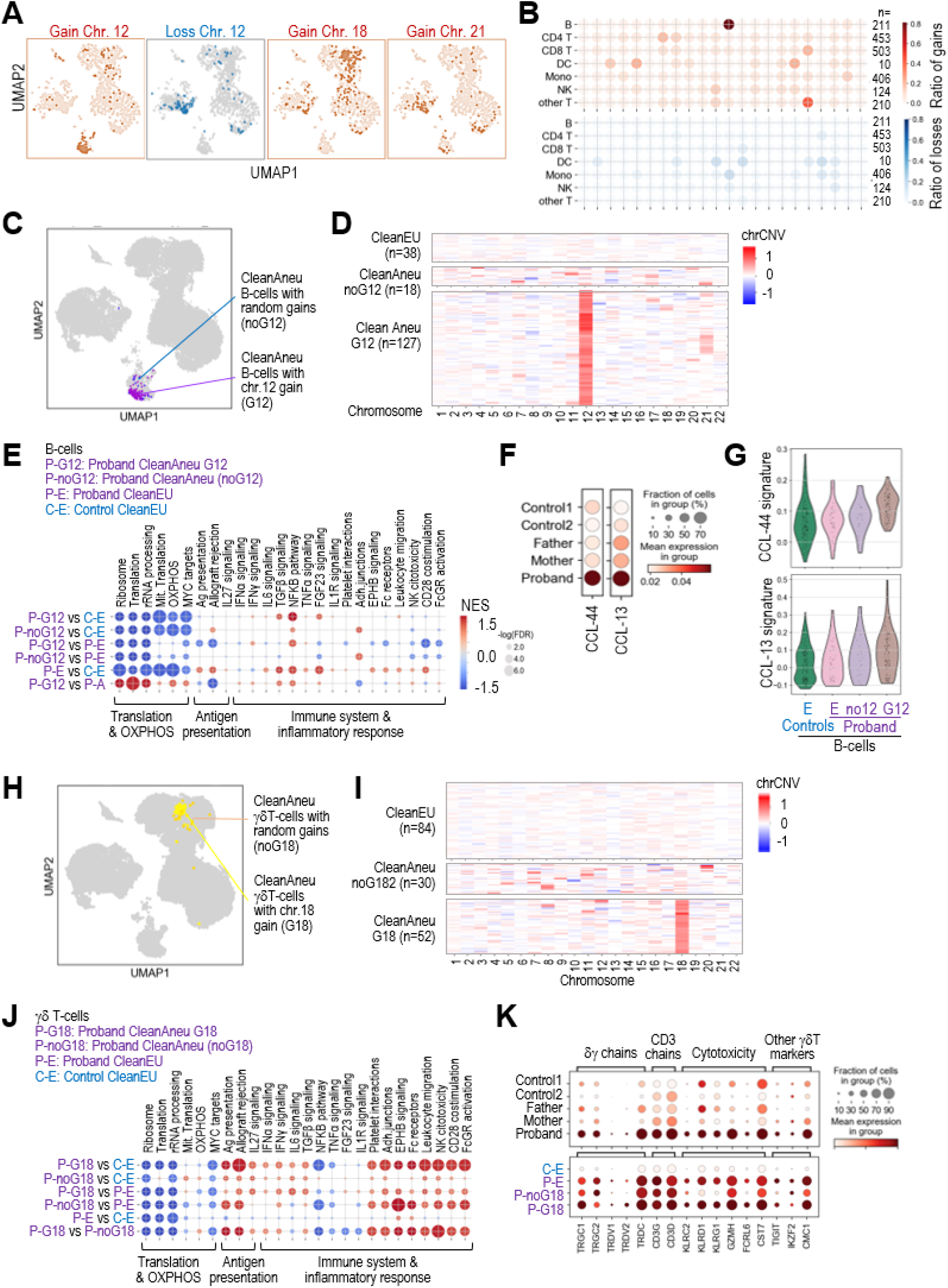
Transcriptomic analysis of random versus clonally selected aneuploidies. **A**, Distribution of aneuploid cells with specific gains (red) or losses (blue) of the indicated chromosomes. **B**, Ratio of cells with gain (red) or loss (blue) of each chromosome among the proband aneuploid cells, showing strong selection of chromosome 12 or 18 gains in B-cells or other T-cells, respectively. The number of cells harboring a specific gain or loss was divided by the total number of aneuploid cells for each specific cell type. **C**, UMAP representation of aneuploid B-cells with gain of chromosome 12 (G12) or random aneuplodies. **D**, Heatmap of the chromosome CNV score (chrCNV) for each chromosome in B cells classified as clean euploid, random aneuploidies with no gain of chromosome 12 (noG12), or aneuplodies with gain of chr.12 (G12). chrCNV values are scaled from −1 (loss of one chromosome) to +1 (gain of a copy). **E**, Gene Set Enrichment Analysis (GSEA) of the pathways selected in Fig. 5 in the indicated comparisons between control euploid B cells (C-E), proband euploid B cells (P-E) and proband aneuploid B cells with gain of chr.12 (P-G12) or random aneuploidies (P-noG12). **F**, Violin and aggregated dot plots showing the expression of a 44- or 13-gene signature typical of chronic lymphocytic leukemia (CLL) in the indicated samples. **G**, Violin plots of the 44- and 13-gene CLL signatures showing enhanced expression in B-cells with chr.12 gains. **H**, UMAP representation of aneuploid γδ T-cells with gain of chromosome 18 (G18) or random aneuplodies (noG18). **I**, Heatmap of the chromosome CNV score (chrCNV) for each chromosome in γδ T-cells classified as clean euploid, random aneuploidies with no gain of chromosome 18 (noG18), or aneuplodies with gain of chr.18 (G18). chrCNV values are scaled from −1 (loss of one chromosome) to +1 (gain of a copy). **J**, GSEA of selected pathways in the indicated comparisons between control euploid γδ T-cells (C-E), proband euploid γδ T-cells (P-E) and proband aneuploid γδ T-cells with gain of chr.12 (P-G12) or random aneuploidies (P-noG12). **K**, Aggregated expression of γδ T-cell markers in the indicated samples with enhanced expression of molecules involved in cytotoxicity in proband γδ T-cells, as well as in γδ T-cells with chromosome 18 gains versus random aneuploidies or euploid γδ T-cells.

Since aneuploidy is frequently detrimental for cell proliferation (1), we next asked whether the clonal selection of chromosome 12 gains in B-cells could provide specific benefits. Transcriptional analysis of proband (P) aneuploid cells with chromosome 12 gains (G12; Fig. 6C,D) versus control euploid cells (C-E) showed inhibition of translation and OXPHOS, similar to other aneuploid cells, but no evidences for immune alteration in these cells (P-G12 vs. C-E; Fig. 6E). Actually, these P-G12 aneuploid cells displayed reduced inflammatory-related pathways when compared to euploid cells from the proband (P-G12 vs. P-E). A direct comparison between proband aneuploid B cells with chr.12 gains versus B cells with random aneuploidies confirmed that G12 cells displayed reduced inflammatory-related pathways (P-G12 vs. P-noG12; Fig. 6E and Fig. S7C,D), suggesting that these clones were protected from the negative effects of aneuploidy. In addition, these B-cells displayed increased expression of two gene signatures that define chronic lymphocytic leukemia (CLL)(*27*) (Fig. 6F). This effect was specifically due to aneuploid cells with chr.12 gains and was not observed in aneuploid B-cells with random aneuploidies (Fig. 6G).

Strikingly, a similar analysis of δγ T-cell clones with chr.18 gains (G18; Fig. 6H,I) indicated that the inflammatory-like response was enhanced in aneuploid γδ T-cells with chr.18 gains (Fig. 6J and Fig. S7E,F). These G18 clones displayed increased expression of cytotoxicity markers such as cell lectin-like receptors (*KLRC, KLRD* and *KLRG* transcripts) as well as granzyme H (*GZMH*), Fc Receptor Like 6 (*FCRL6*) or Cystatin F (CST7)(Fig. 6K). Importantly, only 30% of the expanded γδ T-cell population was aneuploid (Fig. 6I and Fig. S7B), suggesting that the G18 clone arose secondary to an expansion of highly cytotoxic γδ T-cells within the immune response against the aneuploidy-associated stress.

## Discussion

Germline aneuploidy syndromes group heterogeneous entities with mutations in genes involved in cell division. Previous studies have identified mutations in four regulators of the SAC, namely *BUB1B* (MVA1; OMIM#257300)(*8*), *CEP57* (MVA2; OMIM #614114)(*28*) *TRIP13* (MVA3; #617598)(*29*), and *BUB1* (*30*), in addition to *CENATAC,* encoding a spliceasome factor involved in the expression of mitotic regulators (*31*). Common phenotypic alterations in these patients include developmental delay, facial dimorphism and intellectual disability, with a more diverse phenotype in other characteristics such as microcephaly or tumor susceptibility (Table 1).

The biallelic loss-of-function mutations in *MAD1L1*, described in this manuscript, result in lack of full-length protein, defective SAC and a wide spectrum of phenotypic alterations. Whether truncated MAD1 peptides are expressed in this patient is unclear at the moment, although the two mutations prevent expression of the C-terminal globular domain of MAD1, known to be critical for its function in the SAC (*20, 21, 32, 33*). Differential characteristics of this patient versus MVA patients are long survival and no intellectual disability among others. Importantly, the patient shows a striking high cancer susceptibility to a wide spectrum of 12 different neoplasias since early age, including five malignancies by the age of 36. Less than 50% of MVA patients develop tumors and the number of neoplasias per patient is typically reduced to one, with only two MVA1 patients reported to develop both rhabdomyosarcoma and Wilms tumors (Table 1)(*34, 35*).

Whereas germline mutations in SAC components are typically linked to specific cognitive and medical features (*11*), the cellular and systemic consequences of chromosomal instability have not been explored in patients. In agreement with previous studies in cellular or mouse models (*36–40*), our single cell transcriptomic analysis of patient cells suggested defects in translation and ribosome biogenesis, along activation of interferon and NFkB signaling. In *Drosophila*, the reduction in ribosomal proteins associated to aneuploidy has been proposed to trigger the clearance of aneuploid cells from tissues through cell competition (*41*), although whether these mechanisms exist in humans remains unclear. Importantly, single cell analysis allows the comparison of aneuploid and euploid cells, after inferring chromosomal gains and losses from gene expression data. Our data suggest that the inflammatory response is present in euploid cells from the individual with aneuploidy, and leads to increased levels of specific cytokines and inflammatory molecules in the blood plasma.

Interestingly, nuclear humanins are among the top transcripts upregulated in the MAD1 patient. Humanins are short mitochondrial and nuclear-encoded peptides released to the extracellular environment upon mitochondrial stress (*24*) (*42*). The upregulation of humanins and mitochondrial genes, likely as a consequence of OXPHOS defects, might suggest the existence of mitochondrial stress or dysfunction in the patient, which could be a secondary effect of the chronic inflammation (*43, 44*). Remarkably, humanins are also overexpressed in fibroblasts from patients with Down (trisomy 21) syndrome (*45, 46*) in which mitochondrial dysfunction is a prominent feature (*47*). The possible value of humanins as a biomarker in genome imbalance conditions, and its role during the response to aneuploidy remain to be explored. Additionally, multiple mitochondrial components have been shown to act as damage-associated molecular patterns (DAMPs)(*48*), suggesting that humanins and other mitochondrial proteins upregulated in the patient could contribute to the systemic inflammatory response observed.

In addition to random aneuploidies, single cell RNA-seq analysis identified small cell populations with clonal or subclonal gains. B-cells with chromosome 12 gains displayed increased expression of genes that characterize malignant CLL cells (*27*), suggesting possible pathogenic properties. Accordingly, trisomy 12 is seen in approximately 16% of cases of chronic lymphocytic leukemia (CLL) and is associated with poor prognosis (*49, 50*). The majority of CLL patients have been shown to carry prediagnostic B-cell clones 6 months to 6 years before the development of clinically recognized leukemia, suggesting the relevance of close selection in this disease (*51*).

On the other hand, the expanded γδ T-cells with chromosome 18 gains display properties of activated cytotoxic cells with increased expression of granzymes, or killer lectin-like receptors such as *KLRG1*, a with T-cell markers commonly upregulated in patients with chronic inflammation (*44, 52*). The expression of some of these molecules is shared with NK cells, which, using in vitro co-culture assays, were previously proposed to mediate the immune response against aneuploidy (*53*). γδ T-cells play a major role in both innate and adaptive immune responses not only against pathogens but also against own tumor and stressed cells (*54, 55*). However, their specific links with aneuploidy have not been established yet. In summary, our study reports *MAD1L1* mutations in a novel syndrome of mosaic aneuploidy accompanied by unprecendent high susceptibility to a wide spectrum of tumors, some of them never described in MVA patients (Table 1). These phenotypes are accompanied by systemic inflammation and clonal expansion of specific cell populations that may require clinical follow-up. Exploiting the enhanced immune response associated to aneuploidy will provide new opportunities for the clinical management of these patients.

## Materials and Methods

### Study design

We obtained peripheral blood samples from the proband, parents and other family members or controls; biopsies of the hyper, hypopigmented and healthy skin of the proband and from her hyperkeratotic areas of the knee. Paraffin-embedded tissues were obtained from intradermal and compound nevi, pilomatrixoma, pleomorphic adenoma and tubular adenoma of the proband, and also from colorectal and endometrial carcinomas of the proband. We also obtained fresh sample of the third miscarriage of the older sister of the proband, and paraffin-ambedded tissue from the maternal breast carcinoma. Informed consent was obtained from all family members and the study was approved by The Committee for Ethical Research of the Hospital Universitario de Fuenlabrada (Madrid, Spain; reference number 18/26).

### Microsatellite instability (MI) and DNA repair defects

For the evaluation of MI, five quasi-monomorphic markers (NR-21, BAT-26, NR-22, NR-24, and BAT-25) were amplified in the patient’s tumor DNA and in a DNA sample from peripheral blood leukocytes by two Multiplex-PCR. Tumor DNA was extracted from a 30 μm thick formalin-fixed, paraffin-embedded tumor sample using Qiagen DNeasy Blood and Tissue Kit (Qiagen GmbH, Hilden, Germany). For DNA sequencing, selected genes were analyzed using the OncoPlus-GeneSGKit® capture kit (Sistemas Genómicos S.L) and the MiSeq^TM^ platform (Illumina). DNA from peripheral blood leukocytes was extracted using the Maxwell® RSC automatic extractor (Promega, Madison, Wisconsin, WI, USA), following manufacturer’s instructions. Each Multiplex-PCR was performed using the Qiagen Multiplex PCR Kit. Forward primers were end-labeled with 6-Carboxyfluorescein (6FAM) and Hexachloro-fluorescein (HEX) fluorescent marker, according to amplification size. Primer sequences and PCR conditions can be provided upon request. The PCR products were separated by size on an ABI 3730XL DNA analyzer (Applied Biosystems, Waltham, MA, USA). The results obtained were analyzed with Peak Scanner™ v1.0 software.

Detection of DNA mismatch repair (MMR) deficiencies by IHC was performed using a Dako Autostainer or a Leica Bond Autostainer. The antibody clones used were ES05 (1:50) for MLH1, FE11 (1:50) for MSH2, clone EP49 (1:50) for MSH6 and clone A16-4 (1:200) for PMS2.

### Exome sequencing

For Whole Exome Sequencing (WES), we used SureSelect Target Enrichment System v6 (Agilent), following manufacturers protocols. Briefly, 3 μg of high quality genomic DNA was fragmented using Covaris technology, DNA fragments were end-repaired and ligated to adapters prior to hybridization. Sequencing was performed using a 100 bp paired-end on a HiSeq 2000 (Illumina, Inc. USA). Reads were mapped against hg19 using BWA and in-house scripts. Germline variant calling was performed by GATK(*56*) and VarScan(*57*) algorithms, whereas Mutect, Strelka and VarScan were used and combined in order to detect somatic variants. Repeated variants in all samples were filtered out as a step of quality control, besides variants with allele frequency ≤0.3 or read depth ≤10 were also discarded. The variants were annotated with an in-house annotation pipeline gathering information from different sources in order to help its interpretation: germline population datasets (gnomAD), variants genetic variation (Cosmic, ICGC), clinical relevance (ClinVar), genotype-phenotype relationships (OMIM), gene databases on oncogenic role (Tumor Suppresor Gene Database (TSGene)(*58–60*), and in-house database on genes involved in hereditary cancer. For prioritization, variants with a minor allele frequency (MAF) ≥0.1% were filtered out. All non-coding, intergenic, intronic, 3′ and 5′ UTR and synonymous non-splice site variants were also removed. Remaining variants were selected according to a recessive mode of inheritance. Thus, we selected genes carrying homozygous or compound heterozygous variants.

### Metaphase spreads

For karyotyping analysis, peripheral blood cells were stimulated with Phytohemagglutinin-L (PHA) and IL2, and exposed to colcemid (0.5 μg/ml, Life Technologies) for 1 h at 37°C to enrich cells at metaphases stage, and harvested routinely. Briefly, cells were pelleted and exposed to hypotonic treatment (75 mM KCl solution), fixed in cold Carnoy’s solution (methanol:acetic acid (3:1)), and spread onto clean glass slides. Classical G-banding protocol was performed treating the slides with trypsin at 37°C and staining the chromosomes using Leishman solution. Metaphase images were digitally acquired with a CCD camera (Photometrics SenSys) connected to a Zeiss Axioplan2 microscope using a 100x objective, and using Metapher 4 software v3.1.125. Finally, chromosomes were manually classified using Ikaros software v5.0 according to the International System for Human Cytogenetic Nomenclature (ISCN 2013).

### Comparative genome hybridization

Comparative genome hybridization was performed using the Agilent SurePrint G3 Human CGH 60 K microarray (Agilent Technologies) spanning the entire human genome at a median resolution of 41 kb. For the hybridization, 500 ng of genomic DNA from the test samples and human reference DNA (Agilent Technologies) were differentially labeled with Cy5-dCTP and Cy3-dCTP by random priming. An Agilent DNA Microarray scanner G2565CA (Agilent Technologies) was used to scan the arrays, and data were extracted and visualized using Feature Extraction software v10.7. Copy number altered regions were detected with Agilent Genomic Workbench (AGW) software v7.0 (Agilent Technologies) using the Aberration Detection Method 2 (ADM-2) algorithm set as 6, with a minimum number of three consecutive probes.

### Fluorescence in situ hybridization (FISH)

FISH analyses were performed according to the manufacturers’ instructions, as described previously(*61*) on peripheral blood cells treated with colcemid for 1 h, exposed to hypotonic treatment (75 mM KCl solution), fixed in cold Carnoy’s solution (methanol:acetic acid (3:1)), and spread onto clean glass slides. After dehydration (70, 80, 100% ethanol), the samples were denatured in the presence of the specific probe at 75°C for 5 min and left overnight for hybridization at 37°C in a hybridizer machine (DAKO). The probes used detect copy number alterations in chromosomes 7 and 8 (KBI-20031; Kreatech), 20 (KBI-10203; Kreatech) and X (KBI-20030; Kreatech). Finally, the samples were washed with 20×SSC buffer with Tween-20 at 63°C, and stained with 0.5 μg/ml 4′,6-diamidino-2-phenylindole (DAPI)/Antifade solution (Palex Medical). FISH images were captured using a Leica DM5500B microscope with a CCD camera (Photometrics SenSys) connected to a PC running the CytoVision software 7.2 image analysis system (Applied Imaging). Images were blinded analyzed to score for number of FISH signals.

### Variant Calling

Variants were annotated with an in-house annotation pipeline gathering information from different sources in order to help its interpretation: germline population datasets (gnomAD), somatic variants genetic variation (Cosmic, ICGC), clinical relevance (ClinVar), genotype-phenotype relationships (OMIM), gene databases on oncogenic role such as Tumor Suppresor Gene Database (TSGene) and an in-house database on genes involved in hereditary cancer prepared from literature searching candidates. Variants with a population minor allele frequency (MAF) higher than 0.1% were filtered out. All non-coding, intergenic, intronic, 3′ and 5′ UTR and synonymous non-splicesite variants were also removed. Remaining variants were studied considering a recessive model of inheritance, thus selecting genes carrying homozygous or compound heterozygous variants. This approach leaded to 12 genes with at least two non-outfiltered suggestive variants (29 remaining variants), with a single highly suggestive candidate gene (*MAD1L1*), included at the TSGene Database and carriying two protein-truncating variants.

### Single-cell DNA sequencing

For single cell DNA sequencing, genomic DNA was amplified with the GenomePlex Single Cell Whole Genome Amplification Kit (Sigma). Amplified DNA was purified, barcoded and pooled, and sequenced on an Illumina Hiseq2000. Sequencing reads were trimmed to 40 nt and aligned to reference sequence hg19/GRCh37 human genome reference using BWA (0.6.1). QDNAseq (v1.30.0) was used to estimate gene copy number using 100-kb bins, and to generate segmentation plots (*62*).

### Analysis of SAC function

Peripheral blood mononuclear cells (PBMCs) were extracted from blood samples by centrifugation on a Ficoll cushion (Cytiva) 30 min at 1800 rpm at RT, and washed several times in PBS. After 30 min of adhesion step at 37 °C in RPMI, non-adherent cells were cultured in RPMI in the presence of Phytohemagglutinin-L (5 μg/ml, Sigma) to induce T lymphocyte proliferation. After 2 days, PHA was washed and IL-2 (50 U/ml, StemCell Technology) was added to the medium and replaced every 2 days for a time period of 5 days.

To study mitotic progression, PBMCs were synchronized in G2 in by overnight incubation with 5 μM RO-3306 (Selleck Chemicals, S7747). Cell cycle arrest was released by washing cells twice in PBS, and 1 μM taxol (Selleck Chemicals, S1150) was subsequently added to arrest cells in mitosis. Flow cytometry analysis was performed on PBMCs fixed with 4% PFA, permeabilized with 0.5% Triton-PBS and stained with anti-MPM2 (Millipore, 05-368). Cells were washed with PBS and incubated with a secondary antibody conjugated to Alexa-488 (Molecular Probes). Secondary antibody was washed with PBS and DNA was stained with DAPI (2 mg/ml) 30 min at 37°C.

### Protein analysis

For immunofluorescence, PBMCs were attached 30 min to coverslips previously treated with 0.01% Poly-L-lysine (Sigma, P6282). Cells were fixed with 4% PFA, permeabilized with 0.05% Triton-PBS and stained overnight with primary antibodies against: phospho-Histone-3 (Cell signalling, 9701), alpha-tubulin (Sigma, T9026), BUBR1 (kindly provided by Stephen Taylor, University of Manchester, UK), or ACA (Antibodies Incorporated, 15-235). Coverslips were washed twice in PBS and incubated with secondary antibodies conjugated to Alexa-488, 555 or 647 (Molecular Probes) and DAPI. Secondary antibodies were washed three times in PBS and coverslips were mounted in Fluoromount (Bionova).

For western blot analysis, cultured PBMCs were lysed in RIPA (25 mM Tris-HCl pH 7.5, 150 mM NaCl, 1% NP-40, 0.5% Na deoxycholate, Complete protease inhibitor cocktail) 15 min at 4°C, centrifuged 10 min at 14000 g and supernatant was saved for analysis of proteins. Protein extracts were quantified with Pierce BCA protein Assay (Thermo Scientific), mixed with loading buffer (350 mM Tris-HCl pH 6.8, 30% glycerol, 10% SDS, 0.6 M DTT, 0.1% bromophenol blue) boiled for 5 min, and subjected to electrophoresis using the standard SDS-PAGE method. Proteins were then transferred to a nitrocellulose membrane (Bio Rad), blocked for 1 hour at RT in TBS 0.1% Tween-20 containing 5% BSA, and incubated ON at 4°C with specific primary antibodies. Membranes were washed 10 min in TBS-Tween and incubated with peroxidase-conjugated secondary antibodies (Dako) for 45 min at RT. Finally, the membranes were washed 5 min 3 times and developed using enhanced chemiluminiscence reagent (Western Lightning Plus-ECL; Perkin Elmer). The following primary antibodies were used: anti-MAD1 (Abcam, ab10691), anti-MAD2 (Abcam, ab10691), anti-CDC27 (Abcam, ab10538), anti-GAPDH (Cell Signaling, 5174).

Cytokines were analyzed from plasma samples using the Proteome Profiler Human Cytokine Array Kit (R&D Systems, ARY005B) following the manufacturer’s instructions. Relative expression levels of detected cytokines were quantified using Fiji (https://imagej.net/software/fiji/).

### Single cell transcriptomics

PBMCs were isolated from blood samples by centrifugation on a Ficoll cushion (Cytiva) 30 min at 1800rpm and washed in PBS. Fc receptors were blocked using Fcr Blocking Reagent (Miltenyi Biotec, 130-059-901) according to manufacter’s instructions. Dead cells were removed with Dead Cell Removal kit (Miltenyi Biotec, 130-090-101). Viability was assessed (>90%) and cells were subjected to single cell isolation. Generation of gel beads in emulsion (GEMs), barcoding, GEM-RT clean-up, cDNA amplification and library construction were all performed as recommended by the manufacturer. Samples were loaded onto a 10x Chromium Single Cell Controller chip B (10x Genomics) for a targeted recovery of ∼10,000 cells per condition, as described in the manufacturer’s protocol (Chromium Single Cell 3’ GEM, Library & Gel Bead Kit v3, PN-1000075).

All the three classifications, L1 (7 cell types), L2 (30) and L3 (57), were used for more general or specific definitions of cell types (https://azimuth.hubmapconsortium.org/references/#Human%20-%20PBMC). Gene enrichment and pathway analysis was performed with Metascape(*63*), GSEA(*64*), Enrichr(*65*) and their python wrap GSEApy (https://gseapy.readthedocs.io/en/latest/). Ligand-receptor interactions were investigated using cellphoneDB(*25*).

### Inference of aneuploidy from single-cell RNA-seq data

The prediction of aneuploidies was performed using inferCNV (*26, 66*) in their R (https://github.com/broadinstitute/infercnv) and python (https://icbi-lab.github.io/infercnvpy/infercnv.html) implementations, using a window of 250 genes and celltypes predicted in the L3 classification in Azimuth. To estimate chromosomal gains and losses in every individual cell, estimated DNA copy number changes from the gene expression data, using matched control cell types for normalization (Fig. S8A,B). To classify every single cell as aneuploid (i.e. harboring at least one chromosomal gain or loss) or euploid (all disomic chromosomes), we first calculared the average CNV score per chromosome for each cell (chrCNV; Fig. S8C), and established a threshold for classifying each chromosome as monosomic, disomic or trisomic based on the kernel distribution of the CNV score in the proband and controls (Fig. S8C). We also used more stringent thresholds (Fig. S8C) to define “clean” disomies (CleanEU) and either mono- or trisomies (CleanAneu) that may allow a better confidence in the transcriptional comparison between these groups Only cells with at least one mono- or trisomy with the stringent thresholds were considered as CleanAneu (13.6% in the proband vs. 4.0% in controls), and only cells with all disomic chromosomes using these stringent conditions were classified as CleanEU.

### Statistical analysis

Statistical comparisons were performed using Prism 8 (GraphPad), using the specific tests indicated in each figure legend. P values were corrected for multiple testing where appropriate using Bonferroni corrections unless otherwise indicated. Genomic analysis was performed using python 3.8, R 4.1.3 and the packages indicated in the corresponding section.

## Acknowledgments

The thank Stephen Taylor (University of Manchester, UK) and Katja Wassmann (CNRS, Paris) for antibodies and suggestions.

## Funding

Spanish Ministry of Science, Juan de la Cierva programme (CVB)

Spanish Ministry of Science and Innovation-Agencia Estatal de Investigación (MCI-AEI), BIO2017-91272-EXP (SRP)

Spanish National Research and Development Plan, ISCIII, and FEDER, PI17/02303, PI20/01837 and DTS19/00111 (SRP)

Spanish National Research and Development Plan, ISCIII, and FEDER, PI21/01641 (RTR)

Fundación Científica de la Asociación Española contra el Cancer, LABAE20049RODR (SRP)

MCI-AEI/FEDER, RTI2018-095582-B-I00 and RED2018-102723-T (MM)

Comunidad de Madrid iLUNG and scCANCER programmes, B2017/BMD-3884 and Y2020/BIO-6519 (MM)

MCI-AEI, Severo Ochoa Center of Excellence, CEX2019-000891-S (SRP, MM, MU)

## Author contributions

Conceptualization: MM, MU

Methodology: CVB, AGC, AHN, MU, SRP, MM, MU

Investigation: CVB, MT, AGC, AHN, DGS, DR, JP, AO, FM, RTR, SRP, MM, MU

Visualization: CVB, ASB, BP, MM, MU

Funding acquisition: JP, SRP, MM, MU

Project administration: MM, MU

Supervision: MM, MU

Writing: CVB, MM, MU

## Data and materials availability

scRNA-seq data are available at the GEO repository under the accession code: GSE197267. R and Python notebooks used in this work are accessible at https://github.com/malumbreslab.

