## Supplementary Material for "Bilallelic germline mutations in *MAD1L1* induce a novel syndrome of aneuploidy with high tumor susceptibility"

**Supplementary Figures**

Fig. S1 to S7

### Supplementary Figures

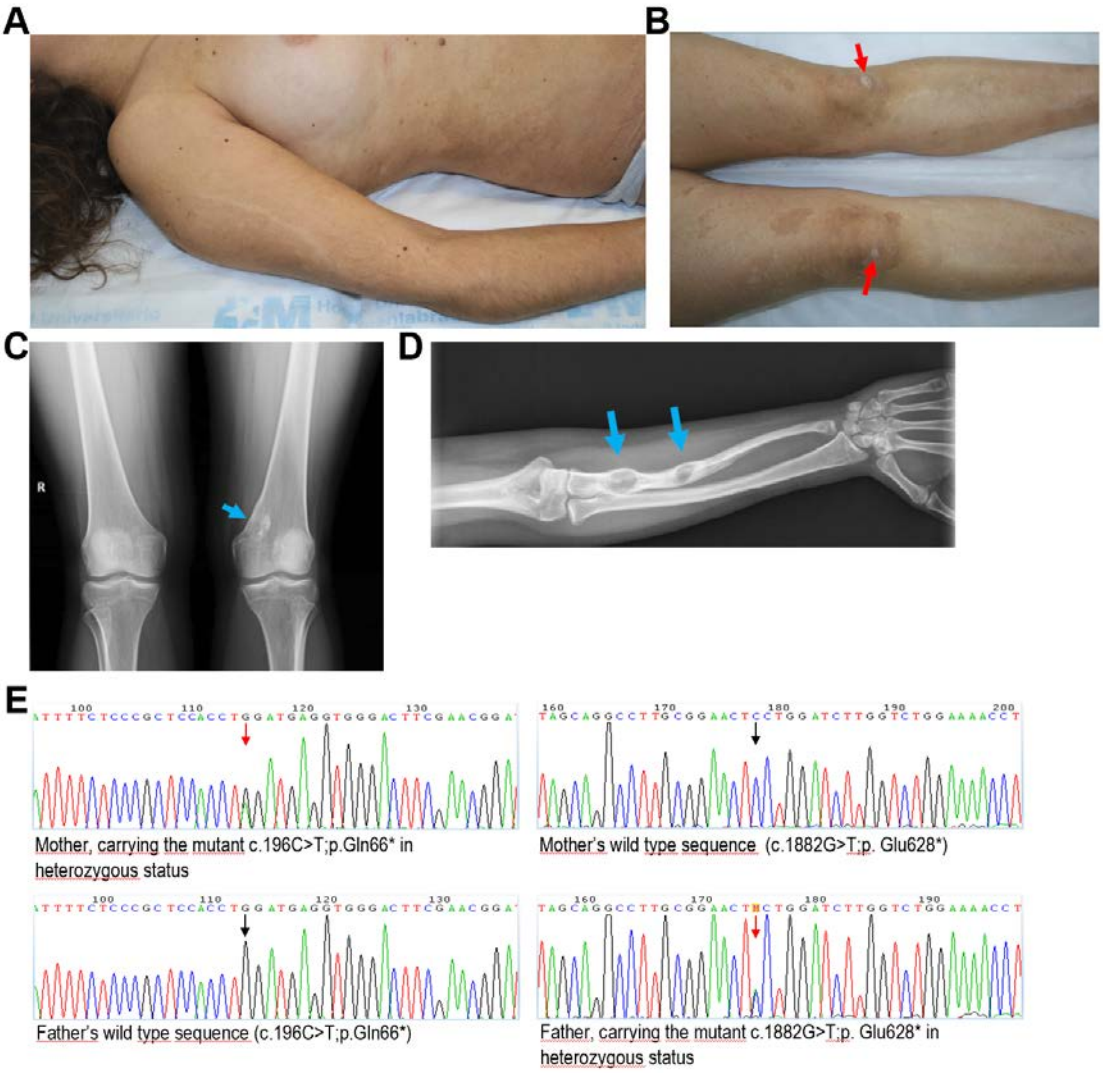


**Fig. S1. Morphological features of the proband and DNA sequencing of family members.** **A**. Blaschkoid hypopigmentation on right arm and elbow deformity. Widespread hyperpigmented lesions of variable tone in the right hypochondrium and lumbar areas. **B**, Multiple café-au-lait spots on both legs with hyperkeratotic plaques with furfuraceous desquamation on both knees (red arrows). **C**, Radiograph of both knees. Lession suggesting enchondroma is evident in left femur (blue arrow). **D**, Radiograph of righ arm demonstrating lessions suggesting enchondromas (blue arrows). **E**, Chromograms showing nucleotide changes in the *MAD1L1* sequence in the parents of the proband. Heterozygous changes are indicated by red arrows.

**A**

| Sample | Chr. 7 Gains (%) | Chr. 7 Losses (%) | Chr. 7 N. Cells | Chr. 8 Gains (%) | Chr. 8 Losses (%) | Chr. 8 N. cells | Chr. 20 Gains (%) | Chr. 20 Losses (%) | Chr. 20 N. cells | Chr. X Gains (%) | Chr. X Losses (%) | Chr. X N. cells |
|---|---|---|---|---|---|---|---|---|---|---|---|---|
| III-5 | 2.9 | 1 | 779 | 5.8 | 0.2 | 779 | 1.9 | 3.7 | 614 | 2.3 | 0 | 731 |
| II-5 | 0.8 | 0 | 605 | 0 | 0 | 605 | 0 | 0.1 | 600 | 0 | 0 | 602 |
| II-6 | 0.3 | 0 | 704 | 0.6 | 0 | 704 | 0.3 | 1.6 | 600 | 0 | 0 | 711 |
| Healthy control | 0 | 0 | 600 | 0 | 0 | 600 | 0 | 0 | 600 | 0 | 0 | 600 |

**B**

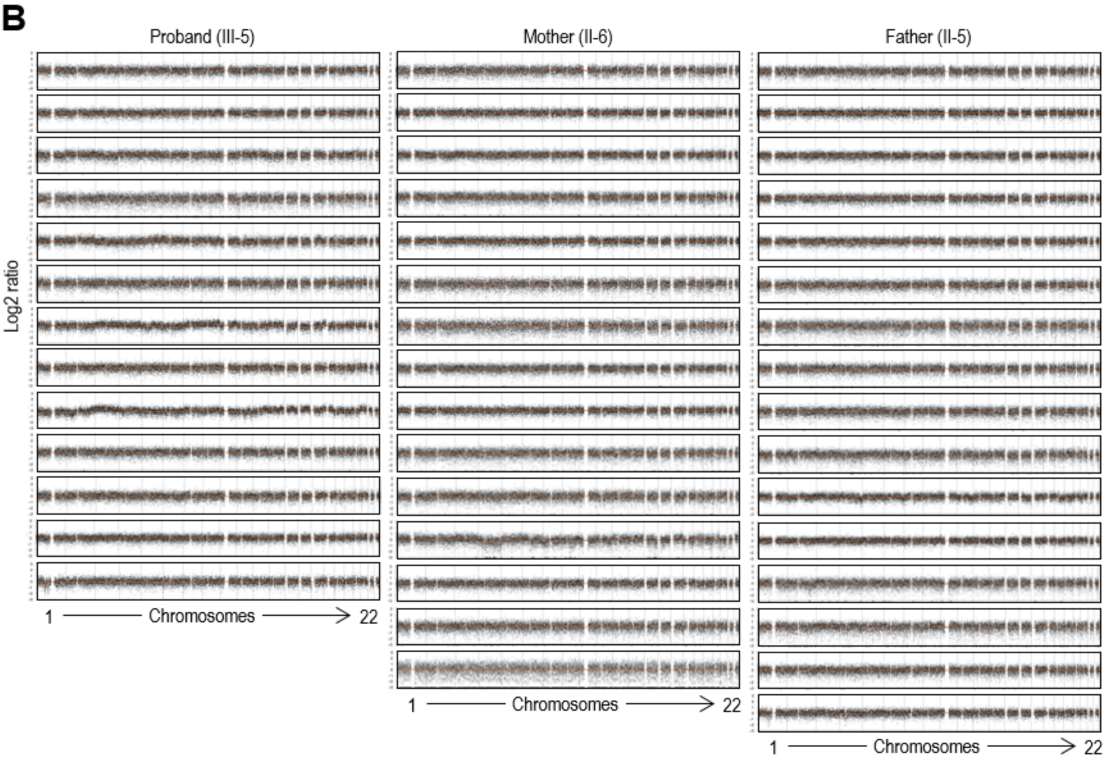


**Figure S2. Analysis of aneuploidy in peripheral blood cells from the proband and controls.** **A**, Quantification of chromosomal gain and losses using two-color FISH with centromeric DNA probes of the indicated chromosomes in blood cells from the proband (III-5), parents (II-5 and II-6) and a healthy control. **B**, Analysis of chromosomal copy number using single-cell DNA sequencing. No significant aneuploidies were found in cells from the father, mother and 13 of the 23 cells analyzed in the proband.

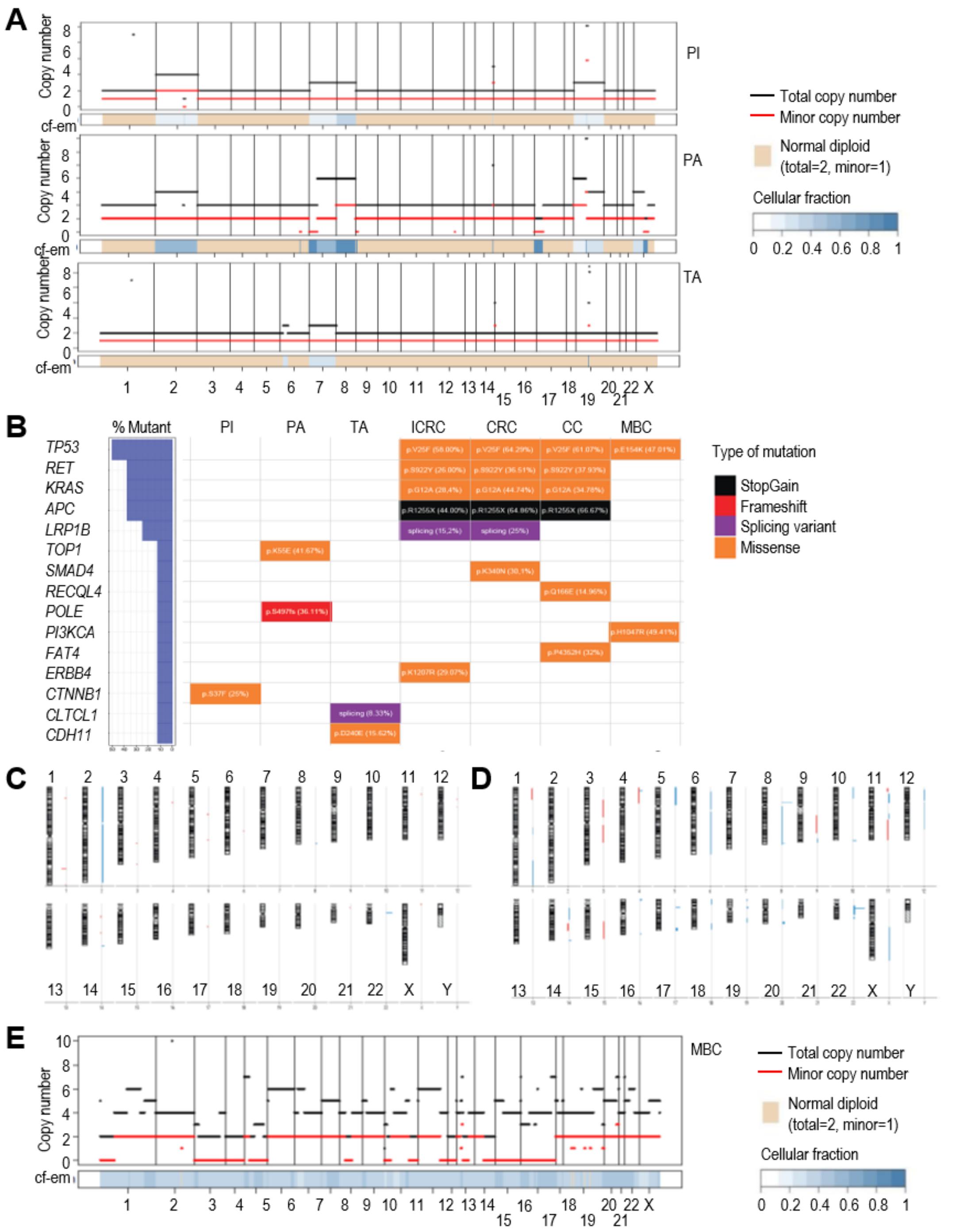


**Fig. S3. Aneuploidy and mutations in pathologies from the proband and relatives.**
**A,** FACET view representing copy number from WES data in different neoplasias of the proband. PI, pilomatrixoma; PA, pleomorphic adenoma; TA, tubular adenoma. **B**, Mutations in cancer-related genes (Cancer Gene Census, https://cancer.sanger.ac.uk/census) after WES. PI, pilomatrixoma; PA, pleomorphic adenoma; TA, tubular adenoma; ICRC, intramucosal colorectal carcinoma; CRC, rectal adenocarcinoma; CC, cervical carcinoma; MBC, metastatic breast cancer in the proband's mother. **C**. Copy number alterations detected by aCGH analysis of an III-4 abortion sample. **D**. Copy number alterations detected by aCGH analysis of an II-6 breast cancer sample. In C,D. blue=copy number gain; red=copy number loss. **E**, FACET view representing copy number from WES data in the metastatic breast cancer (MBC) detected in the proband's mother.

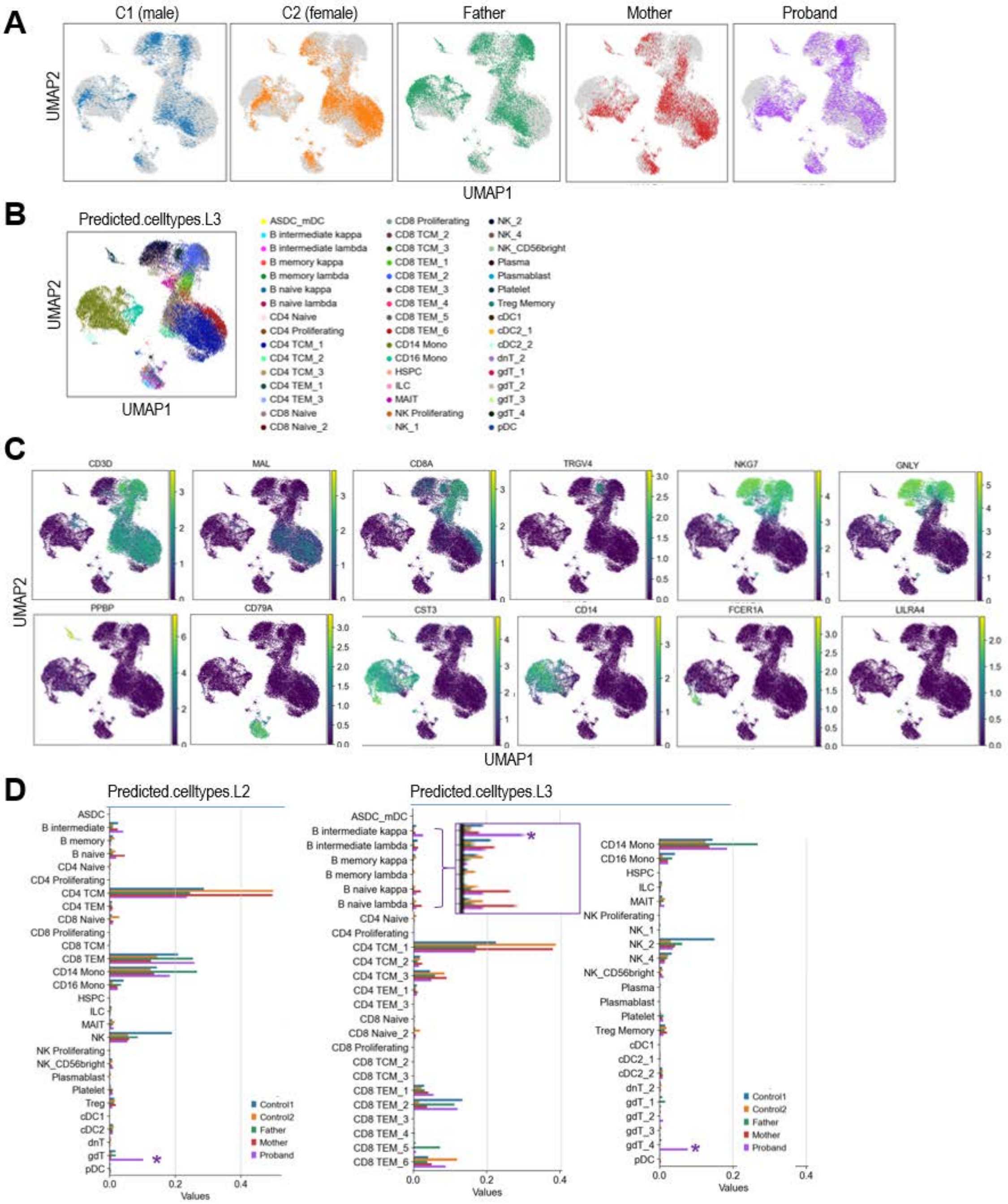


**Fig. S4. Single-cell RNA-seq analysis of peripheral blood mononucleated cells.** **A**, UMAP representation of the samples from the different individuals. **B**, UMAP representation of the L3 classification of cell types (predicted.celltypes.l3 from Azimuth; https://azimuth.hubmapconsortium.org/). **C**, Expression of selected markers of T-cells (*CD3D*), CD4 T-cells (*MAL*), CD8 T-cells (*CD8A*), γδT-cells (*TRGV4*), NK and γδT-cells (*NKG7* and *GNLY*), platelets (*PPBP*), B-cells (*CD79A*), myeloid cells (*CST3*), CD14 monocytes (*CD14*), conventional dendritic cells (*FCER1A*) and plasmacitoid dendritic cells (*LILRA4*). **D**, Percentage of different cell types in the classifications L2 (left) and L3 (right). Asterisks show populations with increased representation in the proband samples.

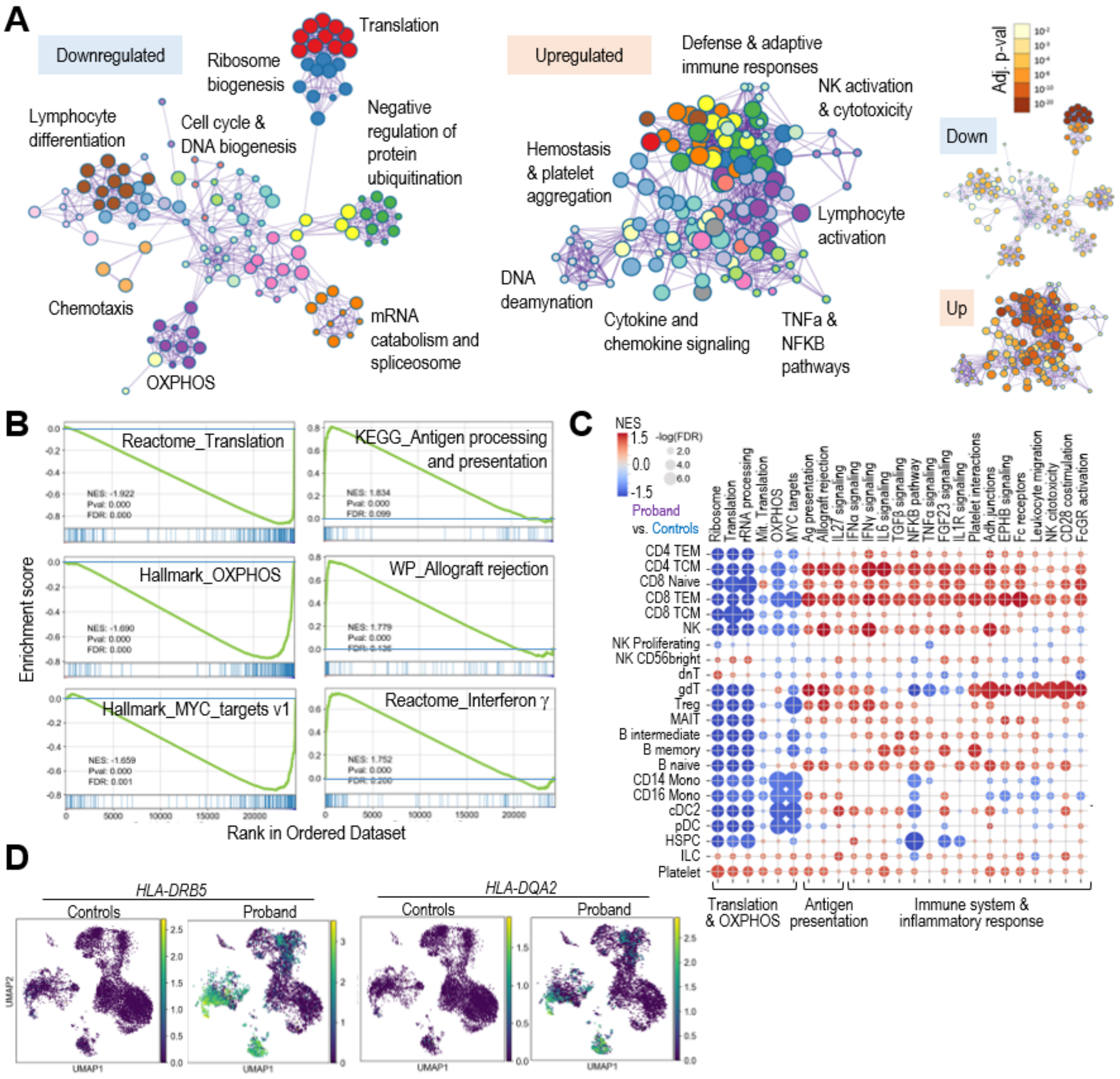


**Fig. S5. Differential gene expression in *MAD1L1* mutant PBMCs. A**, Main pathways down- and up-regulated in from the proband. Pathways were extracted from Metascape using genes down- (LFC <-0.5, adj. pval<0.05) or up- (LFC>0.5; adj. pval<0.05) regulated in proband versus cells from the controls (Control1 and Control2). A general description of the pathways represented in each cloud is shown. The small plots to the right show the adjusted p-value for each specific pathway. **B**, Representative Gene Set Enrichment Analysis (GSEA) of pathways downregulated (left column) or upregulated (right) in proband cells. **C**, GSEA plots used pre-ranked differential expression of genes in proband versus controls in each of the different cell types of the L2 classification. The list of pathways represents a selection of differentially regulated pathways in the different conditions as in Fig. 5, **D**, UMAP representation of the expression of the indicated MHC Class II transcripts in controls (Controls1 + Control2) or proband cells. The distribution of the L1 celltypes is shown to the right as a reference.

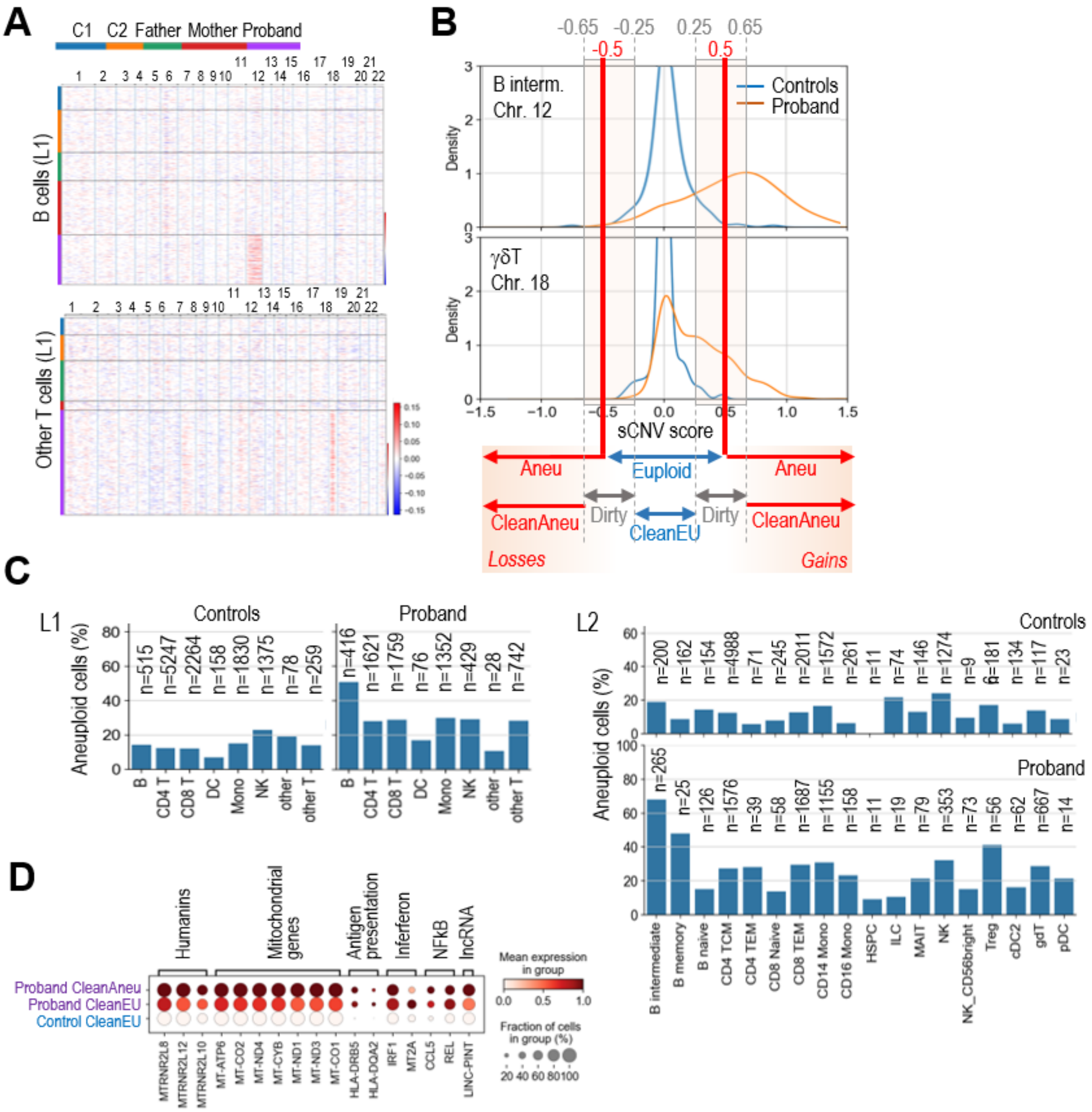


**Fig. S6. Quantification of aneuploidies in scRNA-seq data. A**, inferCNV plots in selected populations from the L1 classification. The scale for the CNV values in each dot of the plot is indicated to the right. Samples are color-coded as indicated. **B**, Schematic representation of the thresholds used to segment chromosomes as monosomic (Aneu with losses) or trisomic (Aneu with gains). chrCNV values were scaled from -1 to 1 using control cells as a reference (sCNV score, scaled chrCNV score). The kernel distribution of the sCNV score for chromosome 12 is shown in B intermediate cells (L2), and for chromosome 18 in gdT cells (L2) from controls (Control1 and Control2) and the proband. -0.5 and 0.5 were selected as thresholds for the initial Euploid vs Aneuploid (Aneu) classification. 0.25 and 0.65 (and their negative values) were used for the classification as clean aneuploid (CleanAneu), clean euploid (CleanEU) or dirty as indicated. Dirty cells were not used for gene expression comparison between CleanEU and CleanAneu cells. **C**, Percentage of aneuploid cells in the different L1 and L2 cell types in control (Control1 and Control2) and proband cells. The number of cells indicated in each group is shown. **D**, Aggregated expression of the set of top genes upregulated in proband vs. control cells (from panel b) in clean aneuploid or euploid cells from the proband, or euploid control cells.

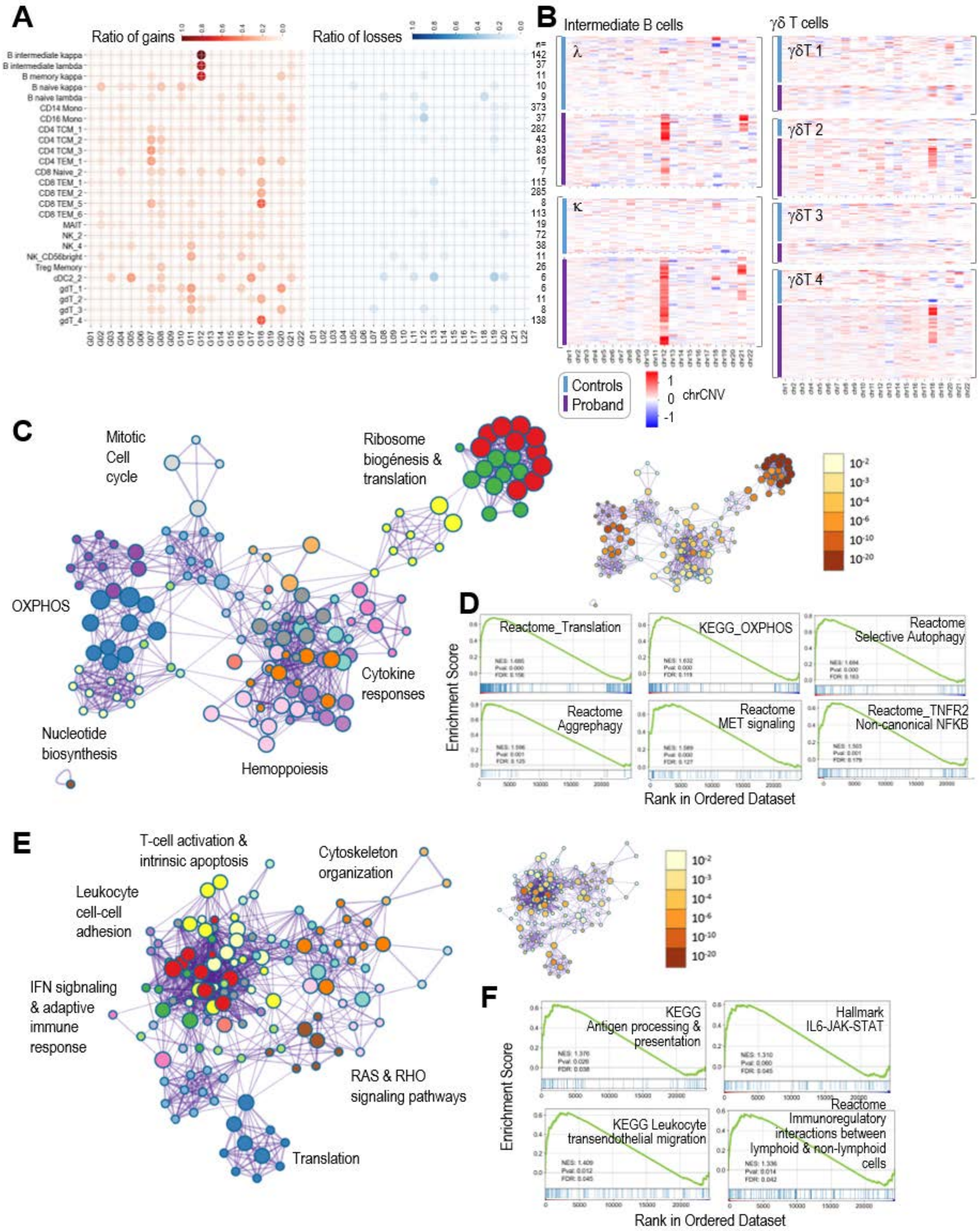


**Fig. S7. Transcriptional analysis of selected versus random aneuploidies.** **A**, Ratio of cells with gain (red) or loss (blue) of each chromosome among the proband aneuploid cells. The number of cells harboring a specific gain or loss was divided by the total number of aneuploid cells for each specific L3 cell type. **B**, Representative chrCNV heatmaps for the indicated celltypes, comparing the values for controls (cells from Control1 and Control2) versus proband cells. The scale bar indicates chromosomal gains (+1, red) or losses (-1, blue). **C**, Main pathways upregulated in proband B cells with gains of chromosome 12 (G12) versus proband B cells with random aneuploidies. Pathways were extracted from Metascape using upregulated genes (LFC>0.5; pval<0.05). A general description of the pathways represented in each cloud is shown. The small plots to the right show the adjusted p-value for each specific pathway. **D**,

Representative GSEA plots of pathways upregulated in proband B cells with gains of chromosome 12 (G12) versus proband B cells with random aneuploidies. **E**, Main pathways upregulated in proband γδ T-cells with gains of chromosome 18 (G18) versus proband γδ T-cells with random aneuploidies. Pathways were extracted from Metascape using upregulated genes (LFC>0.5; pval<0.05). A general description of the pathways represented in each cloud is shown. The small plots to the right show the adjusted p-value for each specific pathway. **F**, Representative GSEA plots of pathways upregulated in proband γδ T-cells with gains of chromosome 18 (G18) versus proband γδ T-cells with random aneuploidies.
